# Network-driven cancer cell avatars for combination discovery and biomarker identification for DNA Damage Response inhibitors

**DOI:** 10.1101/2023.11.20.567526

**Authors:** Orsolya Papp, Viktória Jordán, Szabolcs Hetey, Róbert Balázs, Árpád Bartha, Nóra N. Ordasi, Sebestyén Kamp, Bálint Farkas, Jay Mettetal, Jonathan R. Dry, Duncan Young, Ben Sidders, Krishna C. Bulusu, Daniel V. Veres

**Affiliations:** Turbine Simulated Cell Technologies, Budapest, Hungary; Oncology Bioscience, Research and Early Development, Oncology R&D, AstraZeneca, Waltham, US; Early Data Science, Oncology Data Science, Oncology R&D, AstraZeneca, Waltham, US; Search and Evaluation, Oncology R&D, AstraZeneca, Cambridge, UK; Early Data Science, Oncology Data Science, Oncology R&D, AstraZeneca, Cambridge, UK

**Author notes:** Corresponding author: D.V.V. These authors contributed equally: Orsolya Papp, Viktoria Jordan. These authors jointly supervised this work: Daniel V. Veres, Krishna C. Bulusu.

## Abstract

Combination therapy is well established as a key intervention strategy for cancer treatment, with the potential to overcome monotherapy resistance and deliver a more durable efficacy. However, given the scale of unexplored potential target space and the resulting combinatorial explosion, identifying efficacious drug combinations is a critical unmet need that is still evolving. In this paper, we demonstrate a network biology-driven, simulation-based solution, the Simulated Cell. Integration of omics data with a curated signaling network enables the accurate and interpretable prediction of 66,348 combination-cell line pairs obtained from a large-scale combinatorial drug sensitivity screen of 684 combinations across 97 cancer cell lines (BAC= 0.62, AUC=0.7). We highlight drug combination pairs that interact with DNA Damage Response pathways and are predicted to be synergistic, and deep network insight to identify biomarkers driving combination synergy. We demonstrate that the cancer cell ‘avatars’ capture the biological complexity of their *in vitro* counterparts, enabling the identification of pathway-level mechanisms of combination benefit to guide clinical translatability.

## Introduction

Genetic and phenotypic heterogeneity within a nest of primary tumor cells and their microenvironment has long been one of the major challenges in designing successful and curative treatment plans for cancer patients. The phenomenon contributes greatly to the rapid development of drug resistance and subsequent disease progression ^1–3^. Selecting the appropriate therapy option by optimizing for personalized biomarker patterns increases the likelihood of efficacy ^4–6^, but most cancers progress and subsequential therapy options are needed to tackle disease progression.

Most aggressive human malignancies share common characteristics like aberrant cell proliferation resulting in high replication stress and functional defects in DNA-damage repair (DDR) pathways ^7,8^. The loss of well-functioning DDR has a crucial involvement in uncontrolled tumor growth, disease progression, and therapeutic responses ^9^. Several studies have proved the therapeutic opportunity of targeting DDR in highly aggressive cancers in the clinic ^10–12^, however, patients show variable and often short-lived benefits.

Drug combinations hold the promise of providing a more durable efficacy on cancer cells while managing additive toxicity by decreasing the dosages of individual therapies ^13^. However, finding the right combination partners in the right therapeutic window with potential translatability to the clinic is challenging, therefore, novel approaches are needed to identify combinations that could be ultimately beneficial for patients ^14,15^. One such example includes linking the synergistic manner of a combination with the molecular features of the given tumor mass ^16,17^. In clinical trials, the molecular mechanism determining synergy is often unrecognized, and combinations are often only effective in a patient subset ^18^.

Computational biology is fundamental in developing biological knowledge that translates into clinical practice by revealing potential biomarkers that can predict clinical outcomes and narrowing down the enormous space of potentially synergistic drug combinations ^8,19,20^. Understanding the mechanism of how drug interventions can modify each other could change the perception and prediction of beneficial therapies. Currently, available drug combination prediction methods range from mathematical predictions based on chemical features to pathway-level assessments of biological relations ^21,22^ but display their own limitations. These limitations include: i) lack of explanatory mechanism, ii) uncomprehensive capture of biological feature space, and iii) uncaptured treatment effect at a pathway level. By integrating complementary data reflecting different levels of biological understanding, it is possible to generate a more scalable system of drug therapy prediction with a more granular view of the underlying mechanism.

Network-based approaches are emerging as powerful tools for studying the complexity of cancerous diseases by observing functional interactions between molecules through an interaction network to aid molecular interpretation of the cellular mechanisms behind synergy. These networks have the power to reveal relationships between individual nodes in a certain molecular pathway but also can decipher a higher-level assembly of pathway cross-talk and interconnected cellular mechanisms ^23,24^.

In this study, we used a solution called the Simulated Cell from Turbine Ltd. ^25^ which simulates the signal propagation in a signaling network customized for cancer cells to generate interpretable synergy predictions for combinations of DDR targeting drugs with other agents. By using a primarily protein-protein interaction-based signaling network augmented by cell-type-specific multi-omics data layers that are thereafter simulated using difference equations akin to ordinary differential equations (ODE), but under discrete time, we were able to examine treatment effect at a pathway level (Methods, Fig. 1).

**Figure 1:**
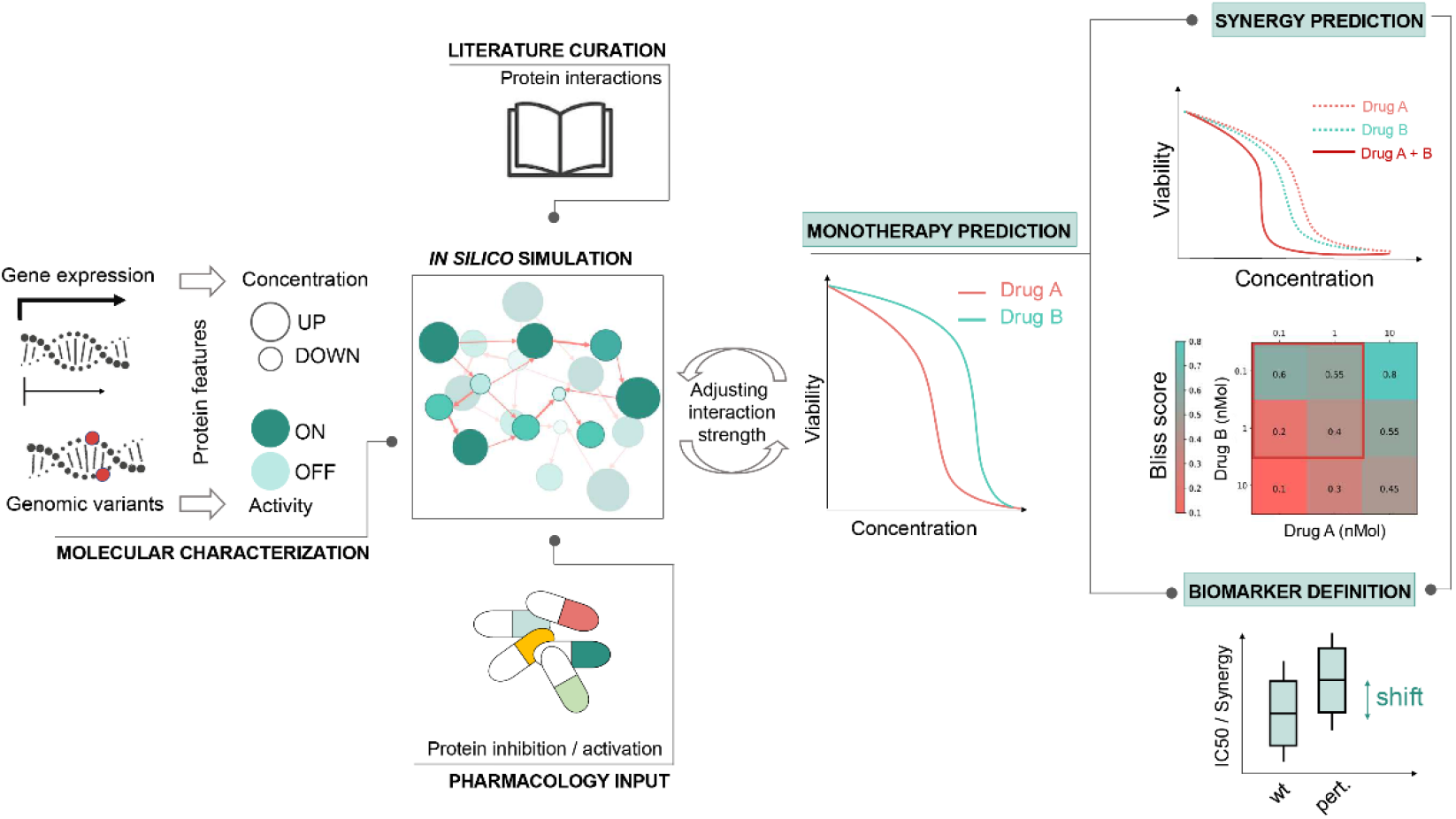
Integration of diverse data types to feed our Simulated Cell to elucidate drug-specific biomarkers as well as to help understand the mechanism of predicted synergy. By using cell line-specific genomic and transcriptomic data, the Simulated Cell can be transformed from a general wiring diagram to network-based *in silico* replica of a cancer cell line. It carries its characteristic mutations beside cell line-specific expression patterns which modify the effect of a node by the binary activity and continuous concentration parameter changes, respectively. Available molecular compound target profiles enable pharmacological perturbation of the signaling network in a dose-dependent and compound-specific manner by modifying the concentration parameter of primary and off-targets of drugs included in the protein-protein interaction network. This approach generates insights into the activity of affected pathways after drug exposure and predicts the cell line’s response to a given intervention by calculating IC50 values and dose-dependent changes in viability. Each Simulated Cell is calibrated to match its *in vitro* counterpart’s IC50 value accurately. This is achieved by setting up the adequate and proportionate contribution of each regulator of a given protein so biological hypotheses are recapitulated on the level of protein interactions that leads to the accurate pathway level signal propagation, and ultimately correct survival response. After the calibration process to establish the network parameters and enable *in silico* experiments, cell line-specific responses of the Simulated Cell to a certain combination therapy–measured by combination synergy and combination-specific viability–are strictly trained on the monotherapeutic effect of the combination partners. By inducing artificial protein alterations, affecting protein activity and/or concentration, combination-specific biomarkers and their effect on combination synergy and cell viability can be identified.

First, we use the *in vitro* monotherapy measurements as benchmarks for the *in silico* cell during manual calibration that aims to match their phenotypes and bring their monotherapy measurements and predictions close to each other. This method enables *de novo* combination synergy prediction. We calculated Bliss independence model-based synergy scores for ∼66,000 cell line-combination pairs. The *in silico* results were analyzed through a transparent benchmarking procedure using experimental endpoints of *in vitro* measurements published in the DREAM Drug Combinations Challenge ^17^.

The Simulated Cell enabled us to interpret the biological rationale determining predictions by following intracellular signal propagation from molecule to molecule, originating from the intervened drug targets to the effector nodes determining cell fate decisions. Furthermore, nodes in the Simulated Cell network could be systematically perturbed in a dose-dependent manner. These perturbations helped the identification of important, non-trivial regulators of the combination-specific response. We used these tools to identify combination-specific biomarkers for poly (ADP-ribose) polymerase (PARP) inhibition combined with ataxia telangiectasia mutated kinase (ATM) inhibition and DNA-dependent protein kinase subunit c (PRKDC) inhibition combined with nuclear factor-kappa B (NFKB) repression while also demonstrating their significant effect on the level of synergy and cell death.

Our findings demonstrate the value of the Simulated Cell as an interpretable network biology-based framework for predicting synergistic drug combination mechanisms. The predictions derived utilizing this toolset could help deliver more precise hypotheses for prospective validation experiments and accelerate the critical translation of *in silico/in vitro* findings to the clinic.

## Results

### Simulated Cell effectively models mechanism of actions for DDR-targeting and some non-DDR-targeting drugs

97 cancer cell lines covering 12 indications, where the *in silico* and the correspondent *in vitro* perturbation results were matching, were used in the project. We supplemented the *in vitro* measurements of the DREAM Challenge ^17^ with further data points from various cellular response databases (Methods; Supplementary Data S1; Supplementary Methods S1) as objective experimental endpoints. Comparing the DREAM data to other sources, we observed overall increased sensitivity, probably due to differences in the length of treatment time (DREAM 120 hours vs. Miscellaneous 72 hours) (Supplementary Fig. S1). The main reason for the observed increased sensitivity is the heterogeneity of the *in vitro* IC50 benchmarking dataset.

**Figure 2:**
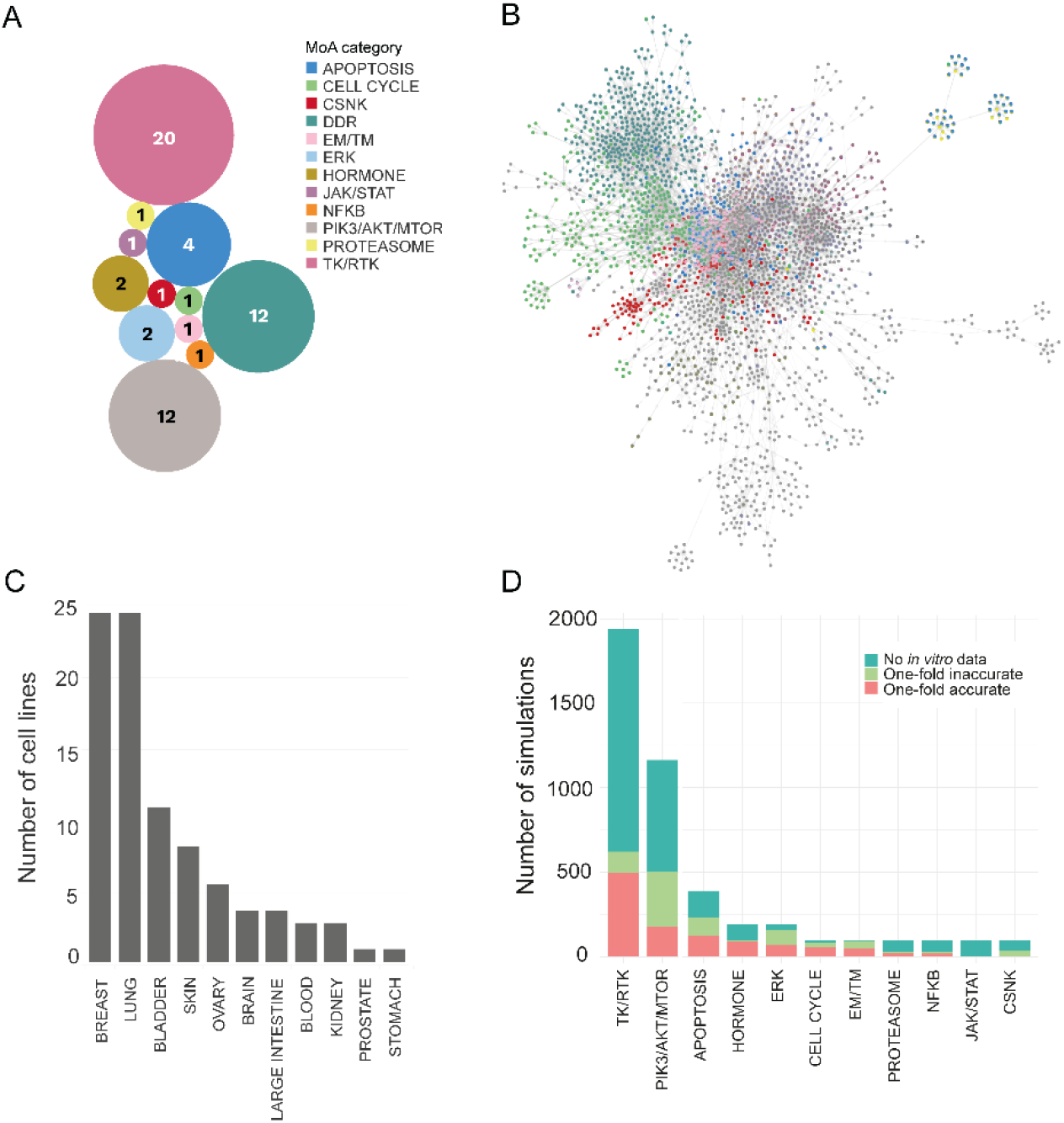
The Simulated Cell encyclopedia and proportions of accurate and inaccurate simulations across the predicted monotherapy landscape. (A) Compound library of the presented combinations. The compounds were categorized into MoA groups based on the pathway membership of primary drug targets with the lowest binding affinity. (B) Snapshot of the network functioning in the Simulated Cell. Our Simulated Cell is based on a graph, where proteins and cellular events are represented as nodes and the physicochemical properties of their interactions are represented as edges between them. DDR pathway members are highlighted in coral, while survival-related pathways, such as different parts of the cell cycle, are colored turquoise. In the model version used for this analysis, our network included 56 modules covering the main cancer-driving pathways with a total of 1997 nodes and 5004 interactions, out of which 14 modules cover the DDR-related mechanisms. (C) Indications covered by cell lines during *in silico* experiments. The cell lines were grouped into indications according to the localization of the primary tumor which the cell line was derived from. (D) Panel represents the number of accurate and inaccurate *in silico* monotherapy predictions compared to *in vitro* measurements in mechanism of action (MoA) categories. The Y-axis shows the overall number of simulations, regardless of the *in vitro* value availability and mechanism of action groups are ranked by the number of one-fold accurate predictions. Unfortunately, due to the low *in vitro* data coverage, there are many cases where benchmarking was not possible. Accurate range was defined as |log10(IC50*in vitro*) – log10(IC50*in silico*)| < 1), meaning the measured *in vitro* IC50 is in the range of (IC50*in silico* / 10, IC50*in silico* * 10) covering an interval of two orders of magnitude. This level of accuracy reflects the variability between two *in vitro* data points in repeated experiments. On the MoA level, the most accurate groups were the TK/RTK and PIK3-AKT-MTOR. Abbreviations: CSNK: Casein kinase, DDR: DNA damage repair, EM/TM: Epigenetic/Transcriptomic modulation, ERK: Extracellular signal-regulated kinase, JAK/STAT: Janus kinase/signal transducers and activators of transcription, NFKB: NF-kappa B, PIK3/AKT/MTOR: Phosphatidylinositol 3’ -kinase(PI3K)-AKT-mTOR, TK/RTK: Tyrosine kinase/Receptor tyrosine kinase.

The drug response of each simulated cell line model was calibrated to reflect their *in vitro* counterparts (Methods; Supplementary Data S1). During the calibration of the Simulated Cell– for the more valuable mechanistic explanations–the priority is to create a correct signaling cascade on a granular, protein level, but only within the limits of keeping an admissible IC50 fit on the majority of the drugs. Due to the size of the network, and within the number of not modeled proteins from the proteome, not all drugs can be fitted *in silico* according to their *in vitro* IC50 values. To test the accuracy of this *in silico* model, receiver operating characteristic (ROC) analysis was applied to the monotherapy response results (Methods). When the results of the individual DDR and non-DDR targeting therapies were aggregated, overall AUC values were 0.7 and 0.47 respectively, with varying performance between individual drugs, especially in the non-DDR category (Supplementary Fig. S2).

### DDR:DDR drug combinations displayed higher synergy than DDR:non-DDR targeting pairs

An *in silico* combinatorial drug sensitivity screen of 684 combinations (12 DDR drugs combined with each other and 46 non-DDR drugs without combining the drugs with themselves) (Fig. 2A) was conducted, each measured in a 16-by-16 dose matrix within a 0-10000 nmol dose range across 97 cancer cell lines (Fig. 2C), resulting in 66,348 combination-cell line pairs measured in total. The generated dataset included *in silico* dose-dependent cell viability measurements and Bliss independence synergy scores ^26^ for all combinations (Methods, Supplementary Methods S2; Supplementary Data S2).

Each cell line-specific combination grid underwent a quality control process to filter out erroneous predictions that do not conform to dose-response principles for one or both compounds (Supplementary Methods S2). Out of the 66,348 cell line-specific combinations, 66,200 combinations remained after quality control and were included in further analytical steps, and only 116 combination grids were discarded due to low quality (Supplementary Fig. S3). After the discretization of Bliss scores, we were able to distinguish between strongly, moderately, and non-synergistic pairwise combinations (Table 1, Benchmarking of the predicted combination synergy scores to experimental data; Supplementary Fig. S4).

**Table 1:** Thresholds of synergy categories determined by K-means clustering. Most of the combination-cell line pairs were not synergistic based on the low *in silico* Bliss values, while the majority of the remaining combinations (20.8% of all) were considered to be strongly synergistic.

|  | **Non-synergistic** | **Moderately synergistic** | **Strongly synergistic** |
| --- | --- | --- | --- |
| **Interval of *in silico* synergy scores** | Bliss score < 0.24 | 0.24 < Bliss score < 0.67 | Bliss score > 0.67 |
| **Number of combination-cell line pairs** | 47583 | 5212 | 13437 |
| **Percentage of combination-cell line pairs** | 71.84% | 7.86% | 20.8% |

The distribution of synergistic and non-synergistic combinations was similar across tumor types. Regarding drug mechanism of action groups, DDR:DDR targeting drug combinations had 2.95-fold higher mean synergy than DDR:non-DDR targeting combinations (0.4944 vs. 0.1672, respectively) (Fig. 3). DDRi:DDRi intrapathway synergy is a logical and known phenomenon. As an example, in BRCA2 mutant high-grade serous ovarian cancer PDX cells, the combinations of PARPi with ATRi or CHKi were synergistic and caused tumor growth suppression and in some cases complete remission ^27^.

**Figure 3:**
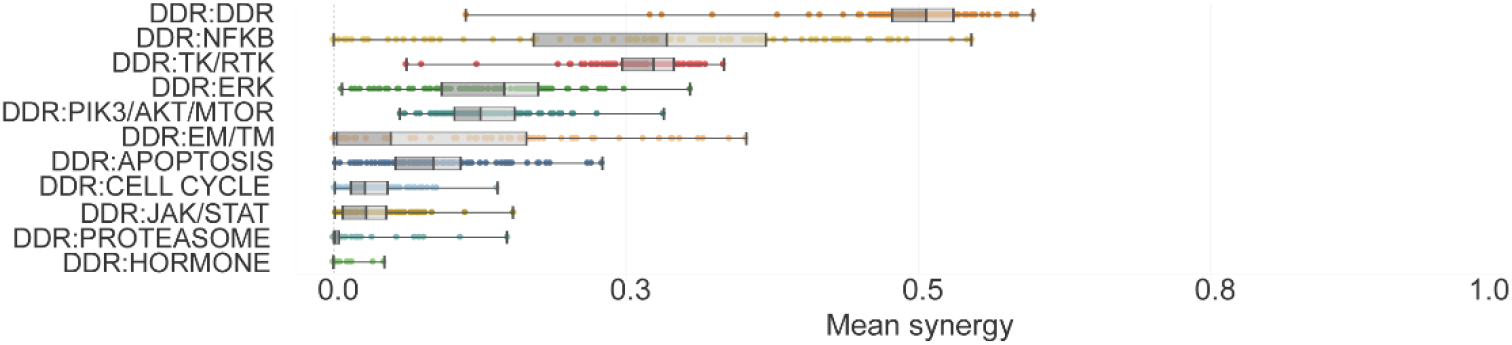
Synergy distribution in different combination MoA group categories. Combination-cell line pairs were categorized into MoA groups based on the molecular targets of the compounds with the lowest binding affinity value. Combination MoA groups were sorted based on the calculated mean synergy. We observed a generally higher synergy in the case of DDRi:DDRi combinations, compared to pairs where DDR inhibition was combined with compounds acting beyond DDR.

To select the most promising combination mechanisms for further detailed evaluation, we prioritized those that we considered truly synergistic, meaning where synergy scores were inversely correlating with cell viability. We calculated killrates for predicting the cell viability decreasing effect of the drugs (killrate = 1 – normalized survival). Some DDR inhibitors combined with compounds acting beyond DDR, were identified as strongly synergistic combinations in specific indications supported by an emphasized cell viability decreasing effect.

Interestingly, from the DDRi:non-DDRi combinations, PRKDCi:NFKBi inhibition performed with a strong synergistic effect in most of the cell lines and indications (Fig. 3, Fig. 4A), and therefore was prioritized for biomarker hypothesis generation analysis (Supplementary Text). Among the better performing DDRi:DDRi combinations, ATM, ATR, CHK inhibitors were observed to be highly synergistic with almost all the other DDR targeting compounds, while interestingly PARP inhibitors were identified as weakly synergistic in combination with other DDR agents (Fig. 4B, Fig. 4C). However, combination of PARPi and ATMi showed relatively strong killrate changes (Supplementary Fig. S5), indicating potential cell killing effect despite the lack of synergy. This observation, together with the clinical relevance of this combination, served as a basis for selecting it for further detailed evaluation.

**Figure 4:**
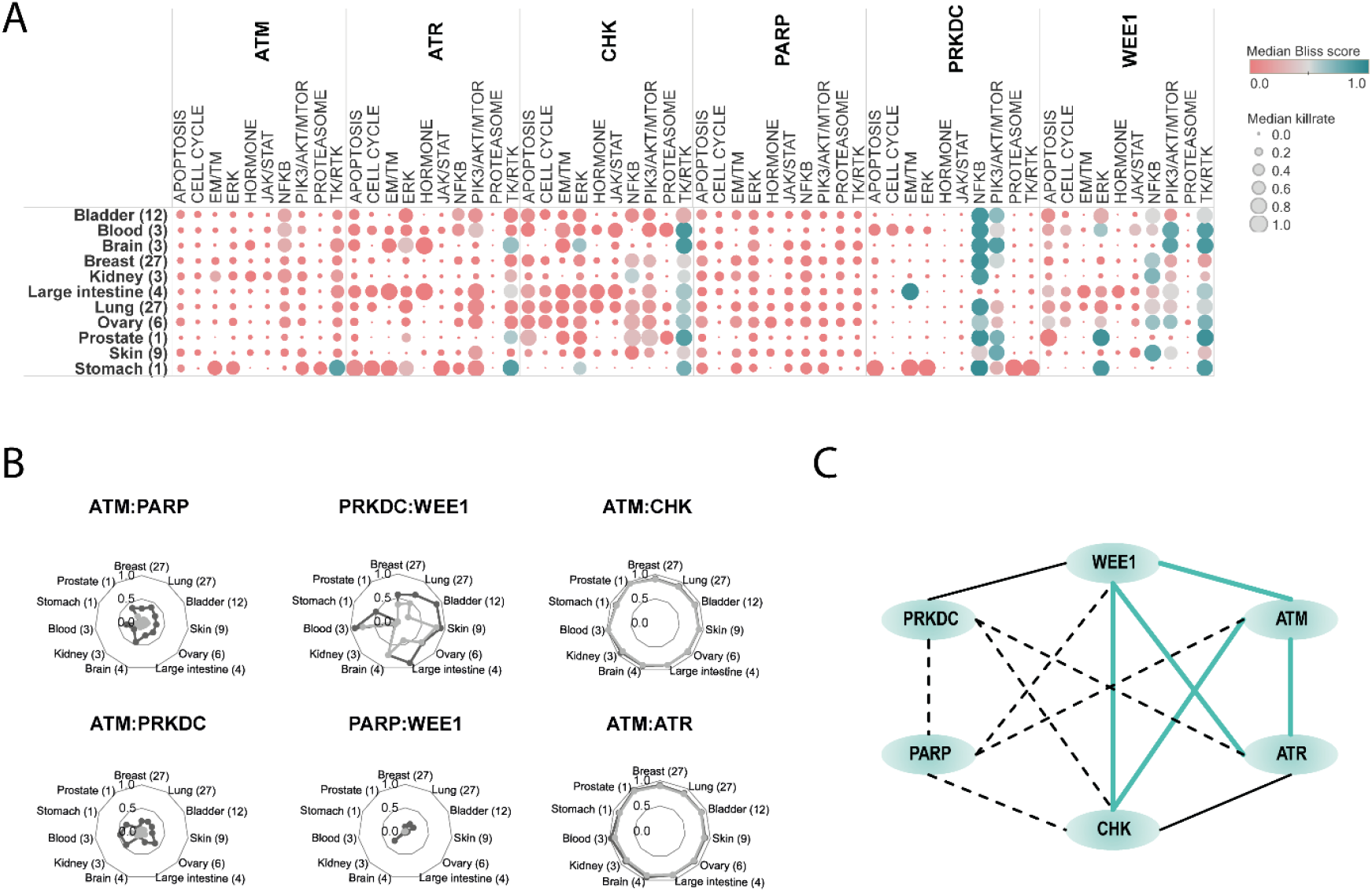
Systematic approach for *in silico* synergy profiling of drugs categorized into different mechanism of action (MoA) groups on cell lines representing various tumor types.. (A) Heatmap for DDRi:non-DDRi combination synergies. Mean of synergy and mean of killrate were calculated for each MoA group for all cell lines in the certain indication group. Combinations that had synergy > 0,5 were highlighted with turquoise color. The size of the circles represents the combination’s effect on cell viability. Bracketed numbers after the name of the indication at the left represent the number of cell lines in that group. The plot shows that CHKi combined with ERKi were strongly synergistic in brain tumor cell lines. PRKDCi represented a strong synergistic effect with EM/TM inhibitors. PRKDCi:NFKBi combinations were highly synergistic and at least cytostatic in many of the indications except for kidney, skin, and colon cancer cell lines. NFKBi performed well in combination with the WEE1 inhibitor only in kidney and melanoma cell lines. Protein synthesis inhibiting drugs (PIK3/AKT/MTORi) combined with WEE1i seems to be strongly synergistic in brain cancer cell lines only. (B) Indication-specific synergistic patterns of DDRi:DDRi combinations. Charts are representing some examples of non-synergistic (ATMi:PARPi, ATMi:PRKDCi, WEE1i:PARPi), moderately synergistic (WEE1i:PRKDCi), and strongly synergistic combinations (ATMi:CHKi, ATMi:ATRi). The latter combinations had a strong cell viability decreasing effect too in all indications, which suggests that ATM inhibitors in combination with ATR and CHK targeting compounds have a cytotoxic effect. Mean of synergy and killrate values were calculated for each MoA group for all cell lines categorized into indication groups. (C) Intra-DDR module combination synergy map reveals frequent intra-DDR module synergy except for PARPi and PRKDCi (except PRKDCi:WEE1i) combinations. Turquoise lines: strongly synergistic relationships between compounds where the cell-killing effect of the combination supported the synergistic phenomenon. Black continuous lines: moderately synergistic combinations, where an intermediate cell viability decrease was observed. Dashed lines: weak or non-synergistic combinations. Abbreviations: CSNK: Casein kinase, DDR: DNA damage repair, EM/TM: Epigenetic/transcriptomic modulation, ERK: Extracellular signal-regulated kinase, JAK/STAT: Janus kinase/signal transducers and activators of transcription, NFKB: NF-kappa B, PIK3/AKT/MTOR: Phosphatidylinositol 3’ -kinase(PI3K)-AKT-mTOR, TK/RTK: Tyrosine kinase/Receptor tyrosine kinase.

### Drug combinations with protein synthesis inhibitors were most accurately predicted with DDR inhibitors

*In vitro* experimental synergy values for our benchmarking analyses were collected from the DREAM challenge published dataset ^17^. Out of the predicted 66,348 combination-cell line pairs, *in vitro* combination synergy values were available for a total of 977 combination-cell line pairs. The coverage of the *in vitro* cell response data differs between compound MoA categories, where availability of benchmarking data is generally superior for DDRi:DDRi over DDRi:non-DDRi combinations (Fig. 5).

**Figure 5:**
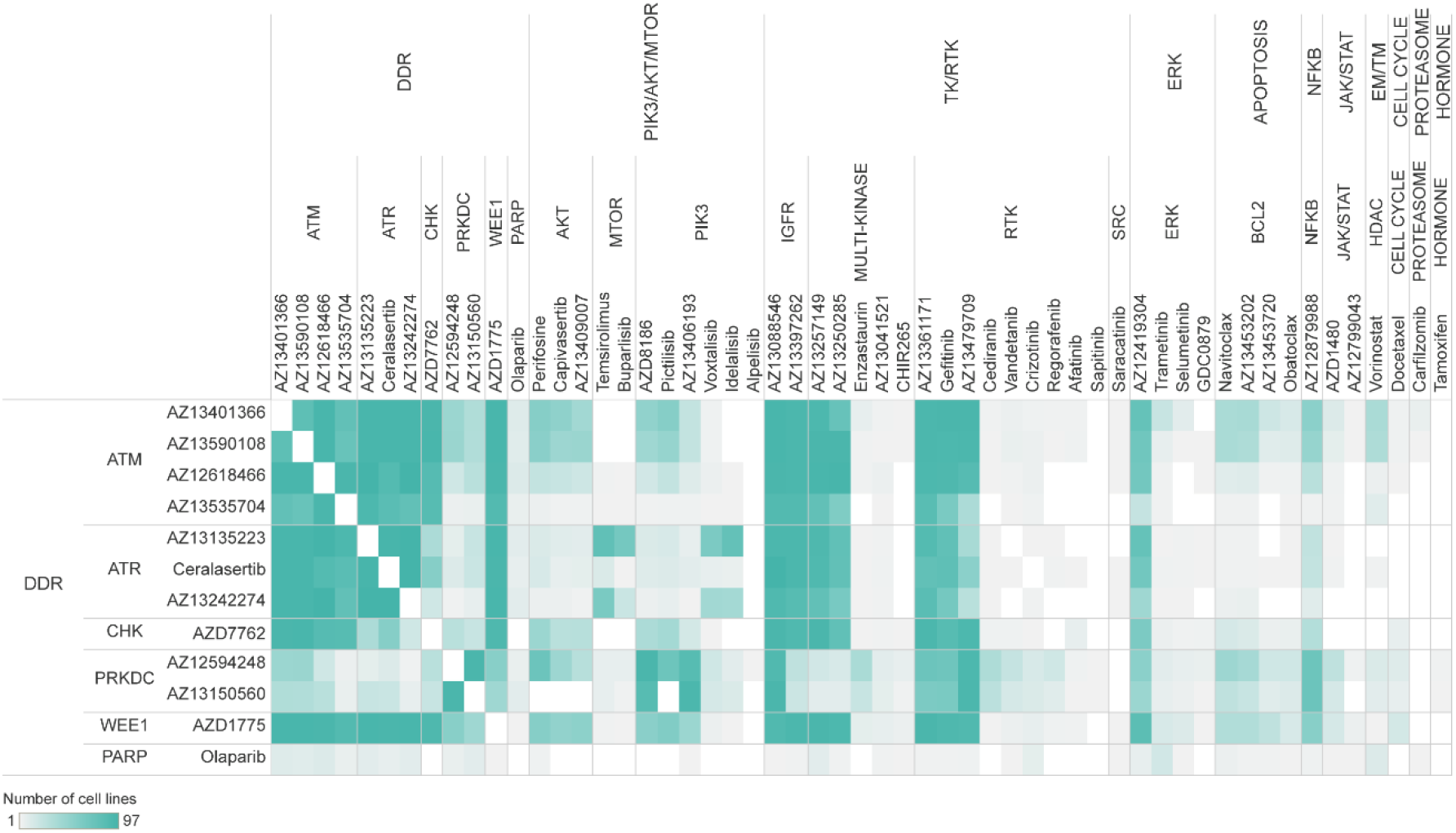
*In vitro* combination synergy measurements used for benchmarking analysis of *in silico* synergy predictions. The heatmap represents the number of cell lines in our *in silico* screen searching for potentially synergistic drug combinations, which are overlapping between the DREAM challenge and our dataset for each exact combination. The drugs were categorized into MoA groups based on the molecular targets of the compounds with the lowest binding affinity value. We had the best coverage of *in vitro* experimental data points in the case of DDRi:DDRi and DDRi:TK/RTKi combinations, also we managed to gather a good amount of data points regarding some DDRi:PIK3/AKT/MTORi combination-cell line pairs. For the rest of the combinations coverage of *in vitro* measurements was poor.

To assess the synergy prediction performance of our *in silico* model, we applied balanced accuracy (BAC) as a metric (Methods). For BAC calculations we used synergy threshold of 20 for the *in silico* results (that would correspond to 0.2 on the original 0 to 1 scale) and 30 for the *in vitro* values from DREAM. For the overall predictivity, we calculated 0.62 for BAC and 0.7 for the AUC from the ROC analysis (Fig. 6A, Supplementary Fig. S6). We observed that drug pairs that were most accurately predicted in combination with DDR inhibitors were mainly protein synthesis inhibitors targeting the PI3K-AKT-MTOR axis, ERK, and other DDR-targeting compounds (Fig. 6B).

**Figure 6:**
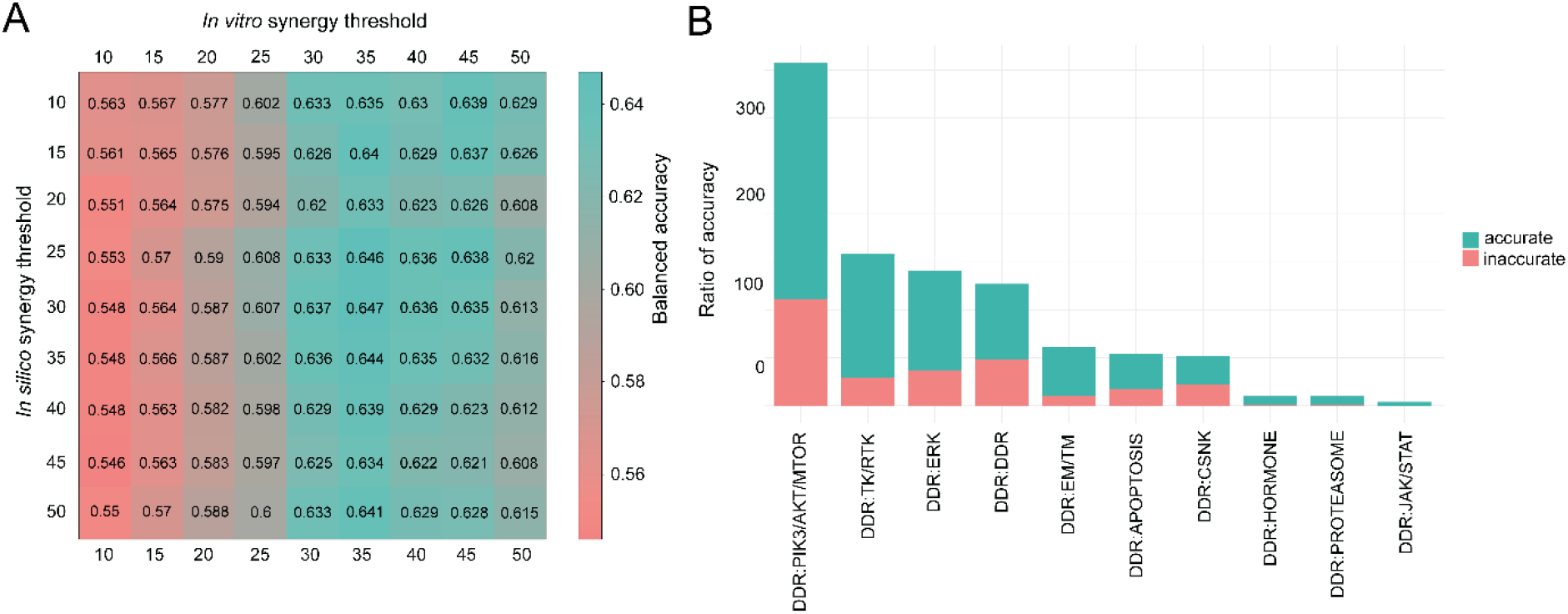
Benchmarking *in silico* combination predictions to *in vitro* synergy measurements. (A) *In silico* balanced accuracy across various synergy thresholds. Matrix represents the threshold value-dependent consensus (expressed in Balanced Accuracy, BA) between *in vitro* and *in silico* synergy. In the DREAM challenge, a threshold of 20 for both *in vitro* and *in silico* synergy was used to distinguish synergistic and non-synergistic combinations. However, the DREAM *in vitro* results were scaled between 0 and 100, while our *in silico* predictions are between 0 and 1, therefore the *in silico* threshold of 0.2 would correspond to it in this case. If we would use this cutoff value for the same endeavor as a gold standard, the balanced accuracy would be just under 0.6 (BAC=0.575, sensitivity=0.37, specificity=0.78). When we set the *in vitro* threshold for synergy to 30 and 20 for *in silico* synergy, our predictive power in balanced accuracy significantly improved (BA=0.62, sensitivity=0.46, specificity=0.78). By increasing the *in vitro* threshold, the number of synergistic data points decreases, and the model was able to correctly detect a bigger proportion of synergistic combinations. This shows that *in silico* predictions are more accurate when stronger synergy is observed *in vitro.* (B) Accurate and inaccurate simulated measurements in different combination MoA groups. Due to the low *in vitro* data coverage, in many cases, benchmarking was not possible. Both *in vitro* and *in silico* synergy thresholds were set at 20, e.g., if both values were bigger or equal to 20, we considered that combination-cell line pair to be accurately synergistic and vice versa. Proportion of accurate and inaccurate simulations were calculated compared to the 977 overall data points. MoA groups are sorted based on the descending proportion of accurate simulations. Abbreviations: CSNK: Casein kinase, DDR: DNA damage repair, EM/TM: Epigenetic/transcriptomic modulation, ERK: Extracellular signal-regulated kinase, JAK/STAT: Janus kinase/signal transducers and activators of transcription, NFKB: NF-kappa B, PIK3/AKT/MTOR: Phosphatidylinositol 3’ -kinase(PI3K)-AKT-mTOR, TK/RTK: Tyrosine kinase/Receptor tyrosine kinase

### Involving more *in vitro* data enhances the predictivity of the Simulated Cell model

It is equally important to understand where the mechanistic model could provide reliable predictions and to identify the governing features of accuracy. In the DREAM challenge, some of the models favored specific subclasses of indications or combination groups. The results revealed that well-predicted classes represented combinations targeting DDR, together with receptor tyrosine kinase pathway and apoptosis inhibitors. We also observed in our analysis that combinations of these mechanisms mainly resulted in accurate predictions.

Our model was calibrated to monotherapy IC50 values for predicting synergy between given compounds and not on combination synergy values (like used in the DREAM Challenge), therefore it is important to evaluate what features drive an accurate synergy prediction. Besides the monotherapy accuracy, cell line coverage and the number of shared targets seemed to be the most important (Supplementary Methods S3, Supplementary Fig. S7-S8), when we evaluated the 977 data points where both *in vitro* and *in silico* synergy values were available. In contrast, when we evaluated the main governing features behind a combination (predicted to be synergistic regardless of this being true *in vitro*), the results revealed the compound-target binding affinities and the monotherapy prediction accuracy for the DDRi compounds as the most relevant features based on all our *in silico* combinations (66,348 data points) (Supplementary Methods S3, Supplementary Fig. S9). These results demonstrate that involving more *in vitro* compound data and increasing the number of elements included in the protein interaction network has the potential to improve both monotherapy and synergy predictivity improvements.

Since our *in silico* signaling network includes several pathway cross-talks, our combination synergy prediction may perform well, even in cases where the monotherapy performance is not accurate. This assumption is supported by plenty of evidence on how drug combination synergies affected by the hindered biological cross-talks of pathways downstream from the primary drug targets ^24,28,29^.

### Molecular alterations that significantly shift synergy are involved in DDR pathways

Many drugs, either as a single agent or as combination therapy, fail in the clinic due to lack of efficacy. Therefore, the identification of biomarkers, predictive of patient response, became an essential part of the drug discovery process, drastically increasing clinical success rates ^30^. We selected two drug combinations, PARPi:ATMi (Olaparib or AZD0156:AZ13535704) and PRKDCi:NFKBi (AZ13150560:AZ12879988) (Supplementary Text) as examples to generate a biomarker hypothesis through a signaling-level understanding of how combination benefit emerges behind the observed synergy and viability score changes.

PARP inhibitors are showing efficacy in the clinic with different biomarker patterns both in monotherapy and in combination. In our dataset, PARP inhibitors were represented with a homogenously poor synergistic effect in combination with other DDR and non-DDR targeting inhibitors. We selected this mechanism as an example to find molecular alterations which tend to shift synergy in a positive manner and have a more prominent cytotoxic effect on cell lines. Additionally, there is preclinical data underlining the potential of the PARP inhibitor Olaparib in combination with one of the ATM inhibitors included in our analysis ^31–33^. Thus, we selected this combination to perform a detailed combination-specific biomarker screen.

Our mechanistic model gives us the ability and flexibility to perturb the activity and concentration parameters of any node included in the signaling network. We simulate the effect of protein over- or under-expression on a given signaling pathway by multiplying or decreasing the concentration of a given node. By knocking it out or making a protein constantly overactivated, we simulate the consequence of hypomorph loss-of-function or hypermorph gain-of-function mutations. These modifications enable us to mimic given extrinsic modifications in a pathway and follow how the signal propagates from one protein to another before and after introducing a particular perturbation. This approach helps identify the biological mechanism behind the observed cell fate after drug exposure in different doses (Methods). After a systematic prescreen for protein alterations causing a combination-specific effect (Supplementary Methods S2, we selected 41 alterations consisting of 33 proteins or protein complexes for the PARPi:ATMi combination to analyze their effect on synergy and cell viability. We further analyzed only those combination-cell line pairs where the dose of the individual combination members at the maximum synergy score was lower compared to the IC50 value of the respective monotherapies, thus exhibiting the potential to decrease drug toxicity and increase drug efficacy. Furthermore, we excluded biomarker-combination-cell line triplets where a significant cell survival decrease was observed but the synergy shifted to a non-synergistic state, as these cases are suspected model artifacts.

**Figure 7:**
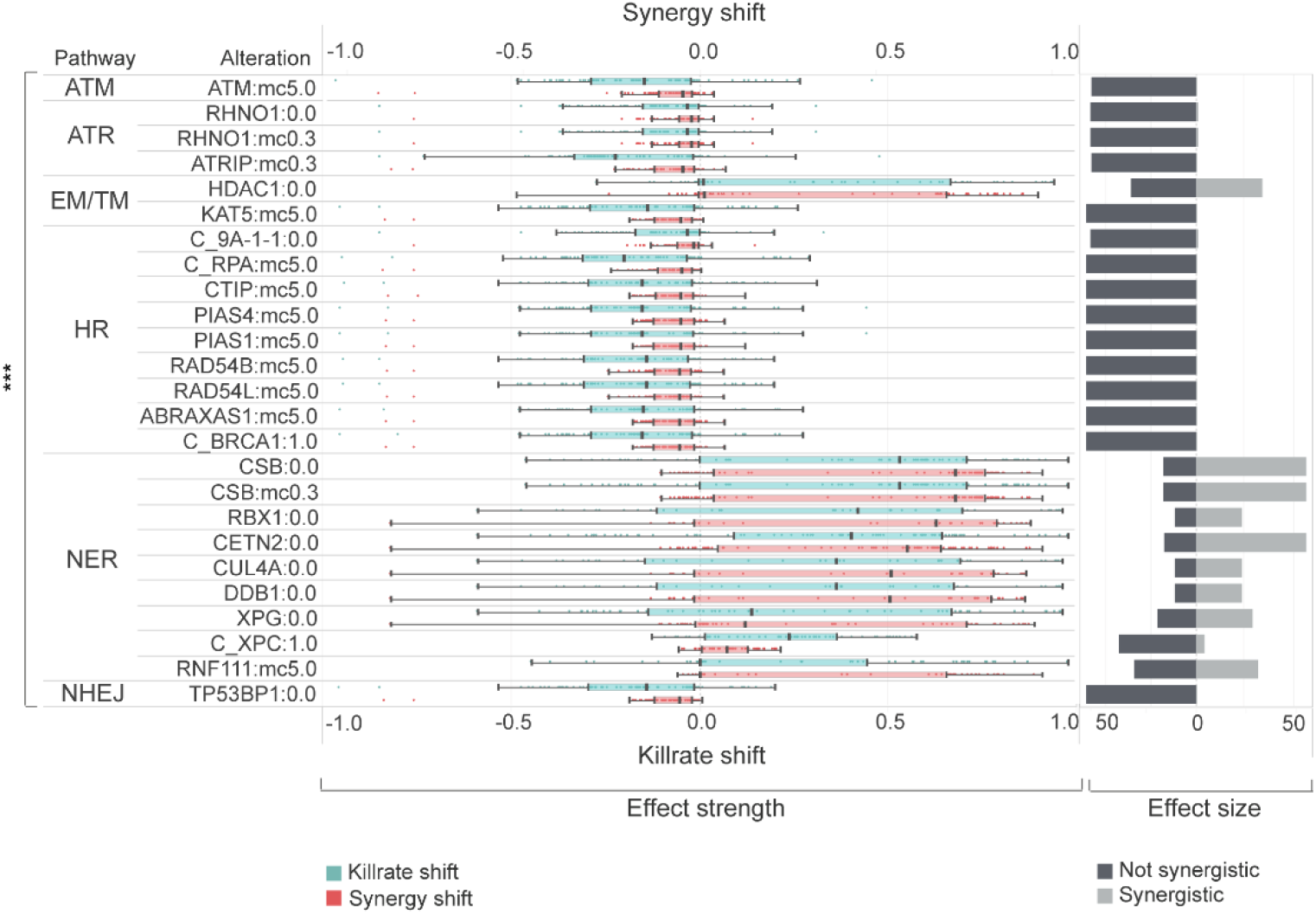
Effect strength and effect size of combination-specific biomarkers for ATMi:PARPi combination. This figure represents the statistically significant (p<0.001, Wilcoxon rank sum test) shift of killrate and synergy influencing biomarkers in cell lines. Alterations for biomarker screening are described in Supplementary M1. Horizontal bars demonstrate the sample size for each biomarker, while the distribution of effect strength components (killrate shift and synergy shift) are represented in boxplots. The loss of NER genes has a favorable influence both on synergy and killrate, while the gaining of HR, ATM, ATR, and NHEJ members have a negative power on effect strength components. Legend: ***statistically significant p≤0.001, 0.0 – loss-of-function, 1.0 – gain-of-function, mc5.0 – overexpression, mc0.3-underexpression, C_ – protein complex. ATM – ataxia-telangiectasia mutated kinase pathway, ATR – ATM and rad3-related pathway, EM/TM – Epigenetic and transcriptomic regulation, HR – Homologous recombination, NER – Nucleotide excision repair, NHEJ – Non-homologous end joining.

We observed that all the alterations resulting in a significant synergy shift are members of different DNA-damage repair pathways. Interestingly, alterations for decreased synergy (potential resistance) were enriched in given DDR pathways, such as homologous recombination (HR) (Fig. 7). In cell lines where HR pathway members were enriched regarding synergy decrease, more than 37% of them carried intrinsic damaging mutations in members of the WNT pathway, causing the pathway’s overactivation (Fig. 8). Many of the most promising alterations increasing synergy were members of nucleotide excision repair (NER), such as CSB, RBX, CETN2, CUL4A, DDB1, and XPG loss-of-function alterations. Inactivation of the epigenetic eraser HDAC1 also led to overactivation, a similar effect as NER pathway perturbation (Fig. 7).

**Figure 8:**
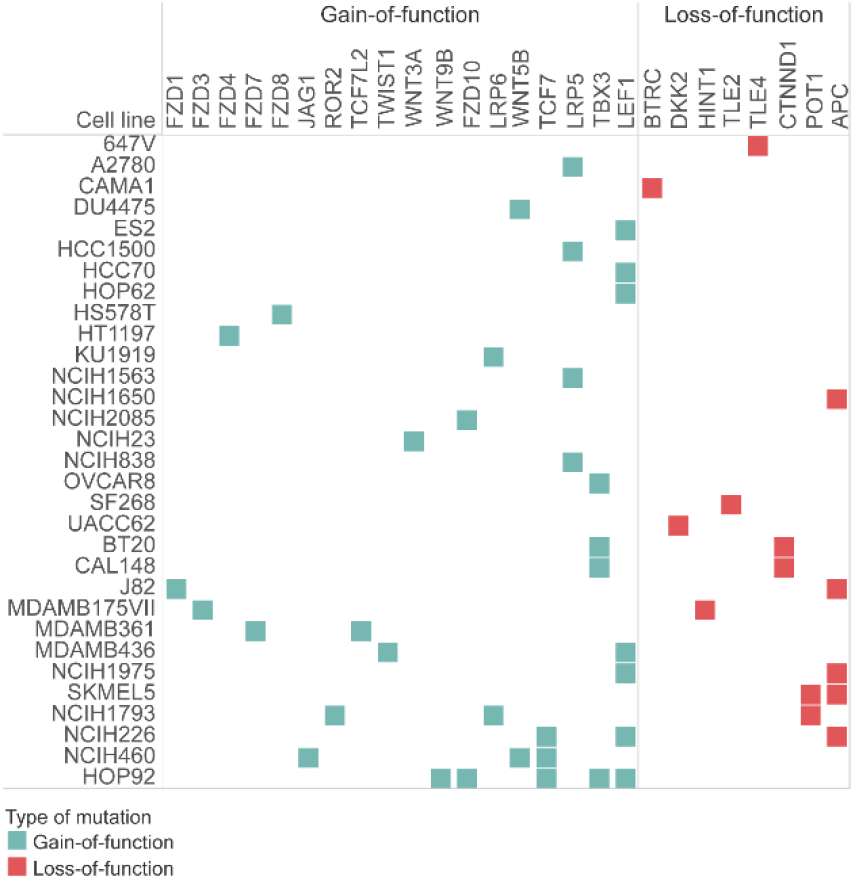
Cell lines, where HR pathway perturbation resulted in synergy and viability decrease had mutations entangled to the WNT pathway. Mutations of cell lines that decreased their viability after artificial perturbations mimicking overexpression and gain-of-function mutations of HR pathway members had a common, intrinsic WNT pathway entanglement. These mutant WNT pathway genes caused the overactivation of the WNT pathway, which is known to be a potential resistance mechanism to PARP inhibition. Gain-of-function mutations appeared in proteins important in activating the WNT pathway, while loss-of-function mutations tended to be appearing in negative regulators of the WNT pathway.

Besides the combination-specific biomarkers, we also analyzed the viability-changing monotherapy-specific biomarkers of the two compounds (Fig. 9, Supplementary Methods S2). In 6 cell lines, 20 molecular alterations were detected as resistance markers for ATMi specifically, while no sensitivity markers were observed. For PARPi, 15 protein alterations were observed as sensitivity markers in 6 cell lines (Fig. 9, Supplementary Data S3).

**Figure 9:**
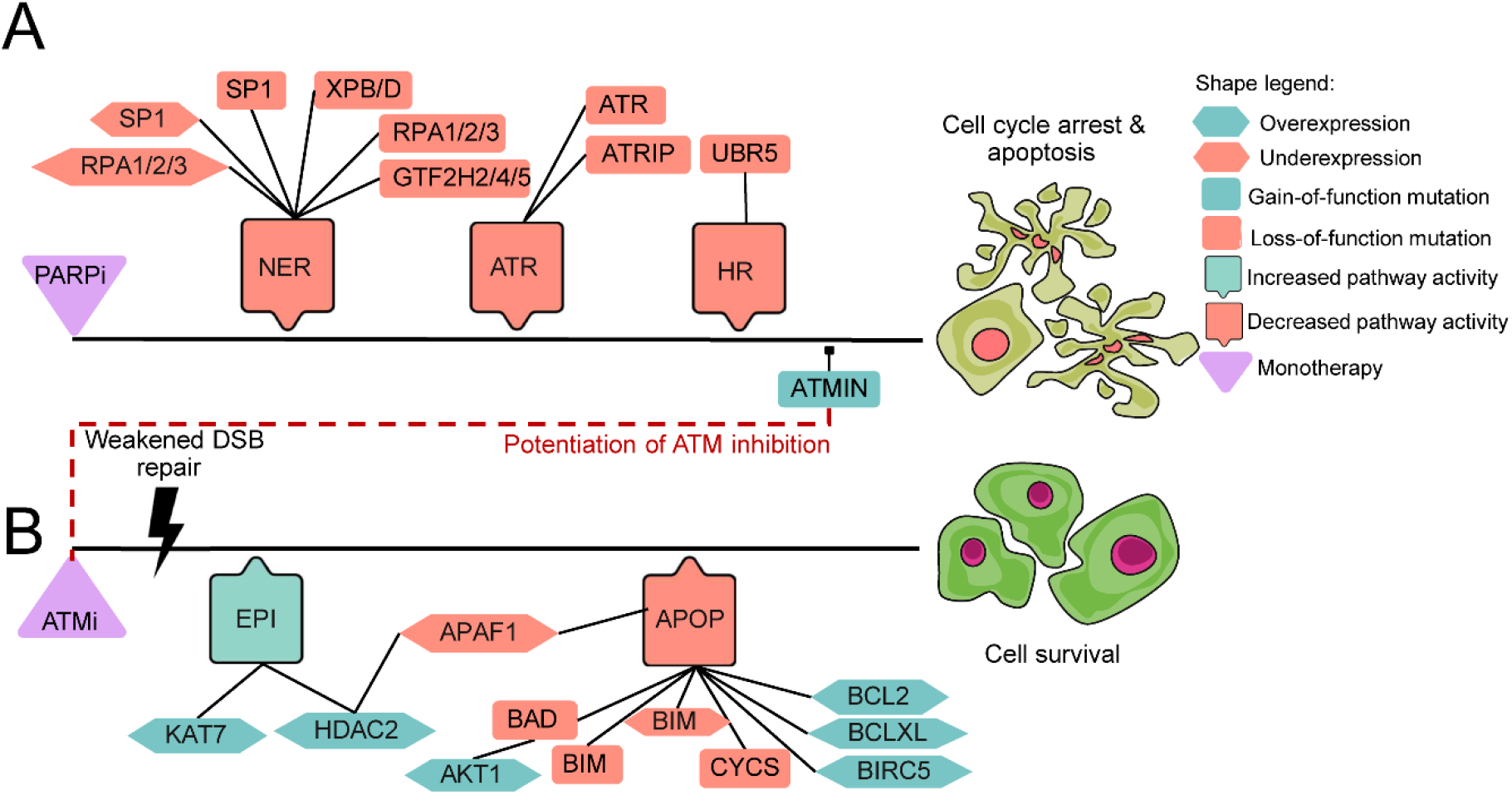
Signaling topology of detected alterations for ATMi and Olaparib can be followed based on the signal propagation in the Simulated Cell network. This visualization represents how each of our predicted monotherapy biomarkers could influence drug responses. Black line represents a theoretical upstream-downstream topology of detected biomarkers based on our signaling network. A) PARP inhibitor-specific sensitizing biomarkers responsible for the downregulation of NER, ATR, and HR pathways which results in triggering cell cycle arrest and apoptosis. B) ATM inhibitor-specific resistance biomarkers are upregulating some members of epigenetic regulation (EPI) which results in the effect of suppressed apoptosis (APOP). Further apoptosis downregulating biomarkers were also observed. Red dashed line represents the point in signaling where the benefit of combining the two chosen drugs could emerge. Since ATMIN gain-of-function surfaced as a sensitizing biomarker of PARP inhibition, this could explain how the combination can overcome the resistance that emerged upon ATM inhibition. The connection between the biomarkers is indirect and shows signal propagation from the drug target to the outcome only.

## Discussion

Our goal was to demonstrate the power of a signaling network-based simulation framework for predicting combination synergy and respective biomarker candidates. Our screen focused on the possible interplay between DDR-targeting drugs with other DDR and non-DDR compounds. We generated combination synergy scores for 684 drug combinations in 97 cell lines by simulating a calibrated signaling network, and without training on the combination data itself. We used the DREAM Challenge experimental dataset to benchmark the accuracy of our drug synergy prediction ^17^. The benchmarking analysis demonstrated that the *in silico* simulation-based results are comparable to the top performers in the challenge (BAC= 0.62, AUC=0.7), even though our model was not trained on any combination dataset before simulations.

Unfortunately, the benchmarking exercise could only be done on the *in silico – in vitro* overlapping dataset of 977 drug combination-cell line pairs, which limits the strength of the comparison. If we break down the non-DDR monotherapy results into MoA classes, among the best performers were drugs targeting tyrosine kinase signaling, the hormone receptor-mediated pathway, and apoptosis, while the worst performers were inhibiting extracellular signal-regulated kinase (ERK) signaling or targets like CSNK (Fig. 2D), indicating variability in predictivity among the different MoA classes. These results suggest that the model’s classification power is generally better regarding DDR inhibitors compared to non-DDR targeting drugs. This observation is in line with the aim of this study to establish accurate modeling of DNA Damage Repair to enable combination synergy and response biomarker predictions. In summary, we demonstrate the potential of the Simulated Cell technology to predict novel combination synergies on a large scale, which combined with the mechanistic explanation behind the observed synergy gives a solid ground for selecting combination treatments for experimental validation.

The benefit of PARPi monotherapy treatment for HR-deficient tumors is already realized in the clinic and approved in high unmet-need populations such as pancreatic or ovarian cancer ^34–36^. However, there is a relatively high chance of drug resistance both intrinsically and acquired ^37^. Understanding the mechanisms behind the emergence of resistance prompts the discovery of novel, more precise biomarkers for patient stratification, and could also lead to innovative combination strategies. ATM is not just a patient stratification marker in prostate cancer for PARPi therapy ^38^, but also a potential target for a combination approach ^31^. However, identifying the precise patient population to target with this DDRi:DDRi combination could be essential in order to limit potential toxicity ^33^.

As combination synergy itself is indicative but not enough to reach combination benefit in the clinic ^14^, our aim was to generate further insight into the combinations by observing the cell-killing effect and hypothesizing biomarkers of increased sensitivity or potential resistance to the drug combination. These results remain to be experimentally validated but the potential of understanding why synergy emerges and how we can use this information for patient selection is demonstrated in two examples.

We hypothesized that combination synergy would appear when PARPi-specific perturbations overcome resistance mechanisms for ATMi (Fig. 9). The loss-of-function of XPB or XPD helicases along with GTF2H2/4/5 could sensitize certain cells to such inhibitors. A loss-of-function mutation of RPA proteins will prolong the initiation of the damage signal by missing the activation of repair inducer ATR at single strand-break sites. Gain-of-function mutations of ATMIN appeared in our prediction because of its ATM inhibiting feature, therefore combining PARP inhibitors with drugs targeting ATM could emerge as a potential combination strategy. ATM inhibition increases the number of double-strand breaks that will grow in a tumor population since the underlying damage response is not initiated, activating apoptotic signaling after a certain limit of errors has been reached. Therefore, underexpression of apoptosis inducers or overexpression of anti-apoptotic proteins could be a potential biomarker. We also predicted that AKT1 upregulation can override the cell viability, decreasing the effect of DNA damage upon ATM inhibition ^39^. Epigenetic apoptosis regulation by APAF1 via HDAC2 or KAT7 could also be a potential biomarker, supported by our predictions as well as literature evidence ^40,41^.

In most of the cell lines treated with the Olaparib:ATMi combination, independent from their tissue of origin, the introduction of deficient NER proteins increased synergy. CSB is considered to be a master regulator of transcription-coupled nucleotide excision repair (TC-NER) as it plays a key role in the recruitment of several proteins to the TC-NER complex ^42^. XPG is another indispensable core protein of the NER machinery, whose function is to cleave the DNA strand at the 3′ side of the DNA damage ^43,44^ and is essential for the 5′ incision by the ERCC1/XPF endonuclease. CUL4A, DDB1, and RBX1 are core components of the cullin-RING-based E3 ubiquitin-protein ligase (CRLs) ^45,46^ complexes which physiologically can promote ubiquitination labeling and proteasomal degradation of target proteins involved in cell cycle progression, transcription, and TC-NER ^47^. CETN2 is a crucial factor in global genome nucleotide excision repair (GG-NER) by acting as an element of the XPC complex ^42,48^. This protein assembly is proposed to be one of the first to bind to DNA damage sites and together with other core recognition factors, XPA, RPA, and the TFIIH complex, is part of the pre-incision complex recognizing single-stranded DNA lesions ^49^. The lack of these crucial components of TC- and GG-NER machinery might be a suitable predisposal factor for impairing single-strand break repair. Since PARP and ATM targeting generate a synergizing mechanism on DSB repair downregulation, increasing the cell’s mutational load will begin apoptosis ^50^.

Beyond identifying NER-related biomarkers shifting cellular response to a more synergistic manner, we also identified the overexpression of HR pathway members (BRCA1 complex, CTIP, RAD54B/L, PIAS1/4, ABRAXAS) leading to decreased efficacy of the combination in cell lines where moderate synergy was detectable. We hypothesize that increased HR activity allows cells to balance between constantly increasing mutational load and decreasing genome integrity, resulting in an increased fitness against the dual intervention of PARPi and ATMi.

Interestingly, the intrinsic mutational profile of these resistant cell lines shared a common feature, namely functionally damaging alterations on the upstream part of WNT signaling (Fig. 8). FZD3/4/10 gain-of-function mutations might cause the WNT ligand receptor, Frizzled, to be constantly activated leading to the inactivation of the destruction complex and subsequent translocation of β-catenin into the nucleus followed by the positive regulation of WNT-dependent transcription ^51^. While FZD3 overexpression was associated with malignancies like lung cancer, FZD4 and FZD10 upregulation was frequently observed in pancreatic and colon cancer, respectively ^52–54^. Interestingly, these resistant cells represented targets (LEF1, TLE2/3) of the WNT pathway being also affected by damaging mutations. LEF1 overactivation serves as a good breeding soil for the upregulation of factors having a crucial role in cell proliferation regulation, like c-MYC and CCND1 ^55^.

WNT upregulation is a known mechanism in drug resistance, especially in the case of PARP inhibitors ^56^. Restored HR pathway activity by for example having function-correcting secondary mutations in BRCA1, managing for the protein to form complexes together with such recombinational well-studied cofactors like the MRN assembly and its further cofactor proteins are well-known resistance biomarkers upon Olaparib treatment, but not reported in combination with ATM inhibition before ^37^.

In order to translate these *in silico* predicted biomarkers to the patient level, first, we estimated the number of patients who could benefit from the Olaparib:ATMi combination therapy. According to our appraisal, more than 120,000 patients with various tumor types could benefit from the combination (Supplementary Data S5). We also analyzed the prevalence of the predicted biomarkers in order to identify segments of the patient population with a higher likelihood of response. Based on the genomic and transcriptomic data of 30 TCGA PanCancer studies, alterations of ATRIP, CUL4A, DDB1, RAD54B/L, and TP53BP1 were the most frequent, indicating their potential to be utilized as patient stratification biomarkers predicting the effectivity of the recommended Olaparib:ATMi combination (Supplementary Data S5).

In order to show that our approach could not just identify already known combinations, but also predicts novel combinations and respective biomarkers, we selected the highly synergistic PRKDCi:NFKBi combination for further evaluation. Whilst there are several reports in the literature of PRKDC and NFKB inhibitor monotherapy activity, we did not find any information about whether these two compounds would potentiate each other’s effectivity or what are the downstream signaling effects of these drugs in combination. We predicted a dozen of potential combination-specific molecular alterations which could shift the synergy and the cell viability beneficially or unfavorably (Supplementary Text, Supplementary Data S4).

In summary, we demonstrate the combination synergy predictivity of our signaling network model-based simulation framework, the Simulated Cell, benchmarking its performance compared to other prediction algorithms in a recent DREAM challenge ^17^. Furthermore, we show that one of the main limitations of such algorithms could be overcome with our model and offer easy-to-interpret mechanistic insights behind the emergence of synergy and ultimately increased cell-killing effect. Based on two examples, the combination of PARPi and ATMi, and the combination of PRKDCi and NFKBi, we were able to not just recapitulate known synergistic effects and underlying molecular features but also predicted novel combinations with a translatability potential based on the identified molecular drivers of synergy.

This model also has its limitations as it requires relatively resource-intensive manual extension of the network, and manual calibration to near each monotherapy *in silico* result to the *in vitro* counterpart. This unfolds mainly in the phenomenon where protein-level biology can be recapitulated *in silico* better than cell survival verdicts. This problem could be solved by the development of a semi-automatic system to extend the network under human supervision and by training the model algorithmically utilizing machine learning. There are also further limitations around the translatability of the results. Limited information is available about the target profile of the drugs, which results in a relatively stronger influence of the primary targets while unknown further polypharmacological effects through potential off-targets are not captured. *In silico* avatars of cancer cell lines are bearing with the general translatability challenges of *in vitro* cancer models, however, the possibility to model the effect of extrinsic molecular alterations which are prevalent in patients is an advantage. Predicted individual transcriptional biomarkers are challenging to translate as these are more dynamic and dependent on environmental factors, therefore complementing the analysis with patient data-based expression analysis is needed, where the data is often limited compared to mutational prevalence.

Overall, we could demonstrate that this framework could be employed for *de novo* combination and biomarker predictions, enabling the selection of experiments to run on a more sophisticated starting hypothesis.

## Methods

### Concept of the Simulated Cell

The Simulated Cell from Turbine Ltd. is a dynamic model of intracellular signaling, integrating a manually curated signaling network with cell line-specific information. Sequenced data on genetic variations can regulate the activity variable of the nodes, and differentially expressed genes influence their concentration. By inhibiting or activating the targets of a test compound according to its binding affinities, the Simulated Cell allows for testing compound efficacy and supplements the results with an understanding of how the biological signal propagates from drug targets across downstream proteins to the final cell fate-determining effector proteins in apoptosis or the cell cycle (Fig. 1) by overlaying sample specific genetic variants and differentially expressed genes as parameters to the interactome. Genomics and transcriptomics data influence the activity and indirectly the concentration of every protein in the signaling network, respectively.

### Structure of the signaling network

The Simulated Cell’s signaling network was created manually by biologists based on trustworthy literature data, that is a common way for creating interactome datasets. Our network has several modules representing pathways (Fig. 2B), where every node, be it an individual protein, complex member, or logical element, has two distinct parameters; activity and concentration. These parameters are influenced by first, the genomic and the transcriptomic data (Fig. 1), and second, the manually set base values for activity and concentration. The directional edges are activity-mediating connections between adjacent nodes that influence the respective activity of the downstream node based on literature data. The effect of the edges along with the manually set base values create logical gates that numerically control whether a downstream node will be activated by its upstream regulators or not (Supplementary Methods S2). The network includes 56 modules covering the main cancer-driving pathways with a total of 1997 nodes and 5004 interactions, out of which 14 modules cover the DNA Damage Response related mechanisms (see Fig. 2B for more details). This connectivity map enables the modeling of module crosstalk and hierarchy in the Simulated Cell.

An ODE network has continuous-valued, continuous-time equations, whereas inside the Simulated Cell, the modified ODE-like model consists of discrete-time, continuous-valued equations. This creates the possibility to model the effect of a protein representing node derived from the binary genomic variations along with the continuous expressional features. The simulation parameters are set up so that after a given number (usually 300) of time steps, 99.9% of the trajectories evolve into a stable attractor state ^57,58^. Attractors are generated by matching specific network outputs with associated phenotypes, such as *apoptosis* and *cell cycle*. Conveniently, these events are driven by particular proteins that can be easily used as the indicator of the alive or dead status of a simulated cell. For example, the DNA fragmentation factor is irreversibly activated when the cell is already dedicated to initiating apoptosis in a specific cellular context and the activity of this protein can be used as the indicator of the activity of apoptosis. The correspondent nodes of apoptosis or cell cycle can reach values between 0 and 1 representing an inhibited (0) or an activated (1) state depending on the upstream signaling events. The Simulated Cell signaling is wired in a way that the cell can never be alive if apoptosis is active, even when its cell cycle is not disrupted. The software predicts a cell to be alive if its readout nodes representing the cell cycle reach at least 60% of maximum effect and apoptotic proteins are not activated at more than 20% of maximum effect.

### Transcriptomics data processing

The Simulated Cell has cell lines from the Cancer Cell Line Encyclopedia (CCLE) in its off-the-shelf library. The transcriptomic input for these cell lines is differential gene expressions between CCLE ^59^ cell lines and non-tumorous tissue data of Human Protein Atlas (HPA) ^60^, a set of 200 samples derived from 32 different (healthy) human tissues. CCLE RNA-Seq data comes from the GDC Legacy Archive, while the HPA data comes from the EMBL-EBI ArrayExpress public repository under the accession ID of E-MTAB-2836 ^61–63^. Thus, the fold changes estimated the relative amount of the given mRNA in each cell line, compared to an average cell used as a baseline, and derived from the healthy expression profiles representing 32 different tissues. At the zeroth step during each simulation, the calculated foldchange value for each protein in each cell line is multiplied by the hand-calibrated base concentration. This way the relative abundance of each protein is represented in the model.

### Determination of genetic variants

In order to use both genomics and transcriptomics from the same source, the Simulated Cell uses genomics from CCLE, in particular from the DepMap (19Q3) mutation dataset ^64^, converted to genome coordinates of GRCh38 by using CrossMap ^65^. Next, its pipeline uses Ensembl Variant Effect Predictor ^66^ (VEP) (we used the parameters listed in Supplementary Methods S1). The Simulated Cell’s input pipelines use this information to map functional consequences to each variant, first using ClinVar (version 2019-07) ^67^, and failing that, predicting using metaSVM and metaLR ensemble scores ^68^ from the commercial version of dbNSFP ^69^ (v3.5) ^67^. In the Simulated Cell, each node is manually annotated as behaving predominantly as a tumor suppressor or an oncogene. Pathogenic mutations hitting oncogenes are assumed to be gain-of-function mutations (unless they are clearly loss-of-function mutations like an early frameshift or stop gain) while pathogenic mutations hitting tumor suppressors are predicted to cause loss of function. The calculated effect of a specific mutation then modifies the achievable minimum or maximum value of the activity parameter of each affected node by either lowering the potential upper threshold to zero in case of loss-of-function mutation or elevating the lower threshold to one if it is a gain-of-function mutation.

### Compound target profile and cell response data

Bioactivity data for the compounds of interest were obtained from ChEMBL ^70^, and additional filtering steps were conducted on the raw data. Firstly, we selected those data points that were derived from *Homo sapiens*. Secondly, we kept data points with confidence scores higher than 5 and consisting of binding assay type. This latter step keeps those data points that provide information about binding affinity. IC50 values above 10.000 nMol were considered ineffective and uncapped data (relation ‘>’) proved to be unreliable for our study, thus, both types were excluded. Additional drug target profiles that were not available in the ChEMBL database and were relevant for this analysis were acquired from AstraZeneca.

*In vitro* compound effects on cell lines have been also received from AstraZeneca. To achieve a greater coverage for the compounds we also downloaded IC50 values from various public drug databases (Supplementary Methods S1). The introduction of a drug onto an *in silico* cell line will result in a lower achievable maximum activity threshold as the concentration of a given drug is increasing.

### Summary of simulation types

The simulations on the Simulated Cell are running in a flexible, easily expandable platform. We summarize the different types of *in silico* experiments, in Table 2. For more details, please see Supplementary Methods S2.

**Table 2.**

| *Experiment* | **Purpose** | **Input parameters** | **Output parameters** |
| --- | --- | --- | --- |
| *Native simulation* | To measure the **native phenotypic behavior** of the cell lines without intervention | – **Node parameters:** base activity, base concentration<br>– **Edge parameters:** direction, type, strength | – Node effects<br>– Attractor scores<br>– *In silico* verdict on cell fate |
| *Monotherapy response screen* | To measure the response of a cell line to a **single drug intervention** | – **Native simulation** input parameters<br>– **Molecular target profile** of compounds | – *In silico* predicted IC50 values of cell lines/drug<br>– Cell viability |
| *Combination therapy screen* | To measure the **synergistic effect of two compounds** in given dose points | – **Monotherapy response** input parameters<br>– **Dose grid** | – *In silico* predicted synergy scores<br>– Doses in synergistic points<br>– Cell viability |
| *Combination specific biomarker screen* | To measure the possible **effect of node parameter alterations on synergy** | – **Combination therapy screen** input parameters<br>– **Altered node parameters:** Under- and overexpression, Loss-and gain-of-function mutations. | – Shift of synergy scores by node parameter perturbations<br>– Shift of cell viability by node parameter perturbations |

The Bliss independence model is a commonly applied statistical model to evaluate compound efficacy in combination through quantification of synergistic/antagonistic effects using the addition law of probability theory (Supplementary Methods S2) ^26,71^. There are multiple reasons why using the Bliss independence model is preferred over other alternative synergy models like applicability on entire dose ranges and in case of non-standard or not available dose-response curves, straightforward probabilistic interpretation ^65,72^. As our main interest was synergy, we modified the Bliss score in a way that we considered every negative Bliss score as zero (total lack of synergism) during the calculation of the aggregated statistics of Bliss scores (Supplementary Methods S2), therefore the probability of antagonism was not assessed in this study.

The benchmarking analysis of *in silico* and *in vitro* monotherapy and combination performances was evaluated by ROC. The predicted Bliss_max_IC50 (Supplementary Methods S2) scores were evaluated in comparison with the *in vitro* Bliss scores. To obtain binary classified *in vitro* measurements, we dichotomized the experimental scores into synergistic and non-synergistic classes using a synergy score threshold of 20. For visualizing and comparing ROC curves the pROC package ^73^ was used in R.

The accuracy of AstraZeneca’s *in vitro* and our *in silico* combination screens were evaluated by a balanced accuracy metric. For the evaluation of combination results, first, an algorithmic method was chosen to discretize the continuous synergy scores. K-means clustering was applied which is a widely used discretization method in biological data analysis ^74–76^.

The statistical significance of the difference between combination synergies aggregated by the compound mechanism group was established by pairwise Wilcoxon rank sum test with Benjamini-Hochberg correction. To test the statistical significance of combination-specific biomarker effects Wilcoxon rank sum test was applied. We concluded statistically significant those effect sizes that got p-value less than 0.001.

### Screening synergy prediction power of the *in silico* model

Synergy scores were binarized for comparison since only Loewe synergy scores were available to be compared to our Bliss independence scores. These two metrics tend to be different depending on the sigmoidicity of the dose-response curves ^77^. Goldoni et al. identified the Hill coefficient as governor of this issue in case of high sigmoidicity, resulting in the overestimation of synergism by the Bliss independence model, while the Loewe additivity model overemphasizes antagonism ^78^. Another difficulty in the comparison is the different calculations of the *in silico* and *in vitro* synergy values. We searched for the maximum value over the whole dose grid, while in the DREAM challenge, the sum of the synergy scores for the dose points is calculated. Based on this information, we applied balanced accuracy (BAC) as a metric to evaluate the synergy predictions. For BAC calculations, we used a synergy threshold of 20 for the *in silico* results (that would correspond to 0.2 on the original 0 to 1 scale) and 30 for the *in vitro* values from DREAM. Regarding the ROC analysis, the same *in vitro* threshold (30) was used.

## Ethical considerations

As the work was completely limited to the use of cancer cell line derived information, which was already published, and all the novel data and insight was generated computationally, ethical approval was not required to carry out the study.

## Data availability

We made all relevant transcriptomic, genomic, and compound targets with cell response data available in the Supplementary Material. The Simulated Cell’s interactome and simulation software is legally protected intellectual property of Turbine Ltd. In case of any further inquiry please contact the corresponding author.

## Code availability

The relevant codes used for data analysis are available in the BitBucket repository: https://bitbucket.org/turbine-public/simcell-combination-synergy/

## Acknowledgments

Authors acknowledge the contribution of the Turbine Ltd. (<u>turbine.ai</u>) team in developing and running the Simulated Cell platform; we especially thank Flóra Bodnár, Mátyás File, Valér Kaszás, Péter Szikora, Szabolcs Komjáthy, and Kristóf Szalay for their contribution to executing the *in silico* experiments and supporting the analysis of the results. The authors also thank the Early Computational Oncology team at AstraZeneca for valuable discussions and feedback. We thank Turbine for funding the project.

## Author information

These authors contributed equally: Orsolya Papp, Viktória Jordán.

These authors jointly supervised this work: Daniel V. Veres, Krishna C. Bulusu.

## Author Contributions

DVV, KCB, JM, JRD, DY and BS designed the study. SK helped to collect the corresponding data from external omics data sources to set the Simulated Cell up. The calibration process was driven by VJ and ÁB. SZH and RB contributed to the statistical evaluation of predictions. BF helped to make the simulation framework accessible for reproducibility check during peer-review. OP performed the analysis of predictions and interpreted the results together with VJ. NNO managed the project team’s work and contributed to the planning and running of the *in silico* experiments beside JV, ÁB, and OP. OP, JV, SZH, RB, NNO, and SK wrote the manuscript. DVV and KCB supervised the project.

