## Supplementary Methods for "Network-driven cancer cell avatars for combination discovery and biomarker identification for DNA Damage Response inhibitors"

Turbine Simulated Cell Technologies

H-1083 Budapest, Szigony u. 26-32

### Supplementary Text

#### Supplementary Text: PRKDCi:NFKBi combination synergy and its monotherapy and combination specific biomarkers

We have selected two drug combinations, Olaparib: AZ13535704 (AZD0156, ATMi) and AZ13150560: AZ12879988 (PRKDCi:NFKBi) as examples to generate a biomarker hypothesis through signaling-level understanding of how combination benefit emerges behind the observed synergy and viability score changes.

There is not much known in the literature nor in clinical trials about how a PRKDCi:NFKBi drug pair would potentiate or antagonize each other's effectivity on modulating cell viability. Our results revealed this combination of mechanisms being highly synergistic and non-synergistic in different indications, therefore both synergy increasing and decreasing biomarkers could be identified in an indication specific manner.

##### Monotherapy biomarkers

We have predicted the sensitizing effect of 34 PRKDCi specific alterations in 12 cell lines, while 14 alteration was observed as viability increasing, resistance causing biomarkers for this compound. In 12 cell lines, in total 81 perturbation were observed as sensitivity markers, while 11 markers were considered to make 8 cell lines resistant to the NFKBi (Supplementary Text Fig. 1).

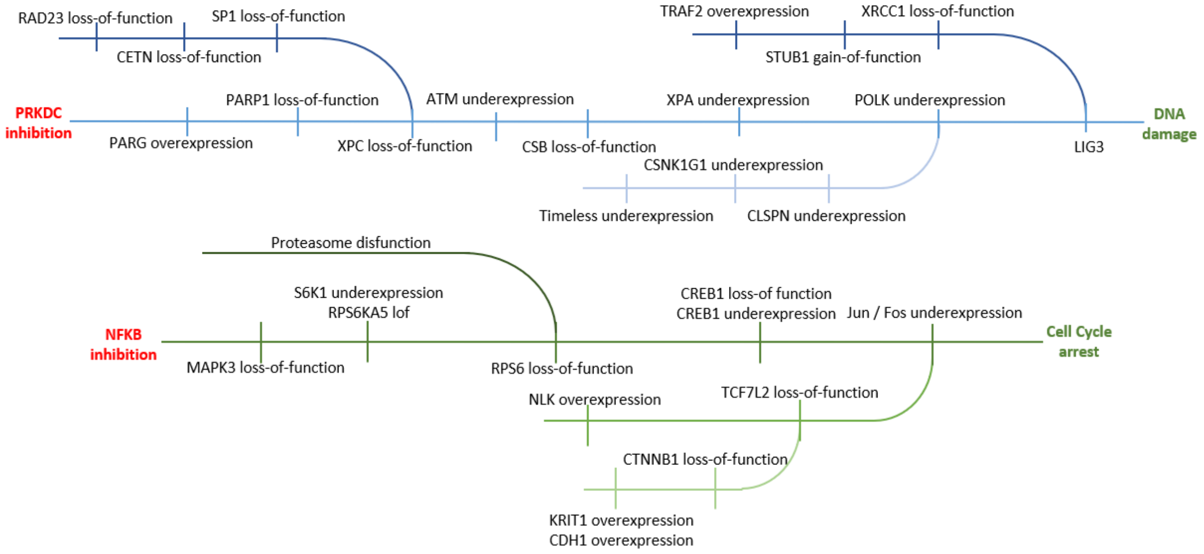

**Supplementary Text Fig. 1. Signaling topology of detected alterations for PRKDCi and NFKBi can be followed based on the signal propagation in the Simulated Cell network**

Visualization of the relationship between the predicted monotherapy sensitivity biomarkers. It represents how they promote and reinforce the cytotoxic effect of both drugs. Loss of the BER or NER damage repair pathway raise the DNA damage load, while many of the mitotic pathway errors lead to cell cycle arrest. The connection between the biomarkers is indirect and shows signal propagation from the drug target to the outcome only.

Inhibition of the NFKB pathway via IKBKB with AZ12879988 will result in the disruption of transcription of a variety of genes working towards cell survival. Our screens showed an array of sensitivity biomarkers upon NFKB inhibition, mostly via inhibiting regulators of protein synthesis and upstream regulators that of, such as RPS6 or its activator kinase RPS6KB1. Parallel to the lack of the synthesis of de novo proteins, a dysfunctional proteasome will push the cell into a stressful state called Unfolded Protein Response (UPR) with a result of an activated apoptosis via the accumulated misfolded proteins during an accelerated protein synthesis need as in the case of cancer. We also predicted the dependency of the transcription factor AP-1 (JUN) and its binding partner c-FOS (FOS) with the consequence of an abrogated cell cycle by lost transcription of its proteins. Our screens also showed some of the upstream regulators of JUN and FOS as important biomarkers by epigenetic regulation of its presence via BARD1 or KAT7. Epigenetic regulation also played a role via the WNT pathway's control of JUN expression. We showed loss-of-function of Catenin β-1 (CTNNB), or the overexpression of its negative regulators such as KRIT1 or CDH1 as sensitivity biomarkers. Same effect was measured downstream of WNT signaling with loss-of-function mutation of TCF7L2 or inhibition of it via the overexpression of its upstream regulator NLK as sensitivity biomarkers in various cell lines.

Abruption of a double-strand break repair pathway such as the non-homologous end joining via inhibition of PRKDC could be the breakdown of the last resort for cells harboring SSB repair related mutations. Disruption of BER pathway via loss-of-function mutation of PARP1 protein or inhibiting it via the constant activation of its counteracting protein PARG represent such case. Another case in our screen was the loss-of-function mutation or underexpression of CSB and underexpression of XPA will inhibit cells to incise problematic nucleotides inserted during synthesis, while underexpression of any subunit of POLK complex or losing a member of the XPC complex will cause issues during the filling of the incised gaps.

Our screens also showed that underexpression of CLSPN (claspin) or loss-of-function mutation of its regulator CSNK1G1 will inhibit POLK to fulfill its role as well. Upon inhibition by any of the drugs losing anti-apoptotic proteins by downregulation will act in favor with the toxic mechanism of given drug, therefore loss-of-function mutation of BCLXL or gain-of-function mutation of upstream negative regulators of BCLXL, such as RXRA, RXRB or HINT1 ended up as sensitivity biomarkers in our screens. Similar to the overall sensitivity of the apoptotic pathway proteins, activating mutation in the RAS-RAF axis such as the RAS-related protein RALB or its GTPase activators RALGAPA1,2 and RALGAPB will end up as resistance markers. Similarity between the importance in apoptotic regulation shows the potential of this combination therapy option.

#### Combination specific biomarkers

There were no biomarker candidates mentioned in the DREAM challenge being specific for this MoA combination. After a systematic prescreen for those protein alterations which are causing a combination specific effect (Supplementary Methods S1), we selected 166 alterations consisting of 118 proteins or protein-complexes for the PRKDCi:NFKBi combination to analyze their effect on synergy and cell viability. We have further investigated only those combination-cell line pairs, where the dose of the individual combination members at the maximum synergy score was lower compared to the IC50 value of the respective monotherapies. Furthermore, we also excluded those biomarker-combination-cell line triplets, where significant cell survival decrease was observed but the synergy shifted to a non-synergistic state, as these cases are suspected model artefacts.

The systematic biomarker prescreen confirmed that several pathways and their members can be relevant regarding shifting both cell viability and synergy (e.g. AKT, JAK-STAT, MAPK, p38-JNK) in case of this combination. Overexpression of STAT3, JUN, FOS increased both synergy and cell killing relative to non-perturbed cell lines. Although the sensitizing effect strength of JUN was statistically significant, its synergy increasing role is considered highly context specific, since JUN itself is transcriptional factor bearing context specific functions. Similarly, the under-expression of STAT3 (p=1.78e-8), overexpression of PPP2CA (p=2.34e-8) or inactivating mutations in ATM or MID1 (p=1.28e-6 and 2.34e-5) were resistance biomarker, since the cell's killrate were lowered by them, thus made the cells less vulnerable to cell death. Synergy was also decreased when their effect were compared to wild-type cell lines. We also observed loss-of-function alterations of members of the mTOR signaling near to the mTOR-apoptosis crosstalk (mTORC2 complex and its complex members, PROTOR and RICTOR) with synergy decreasing effects (Supplementary Text Fig. 2).

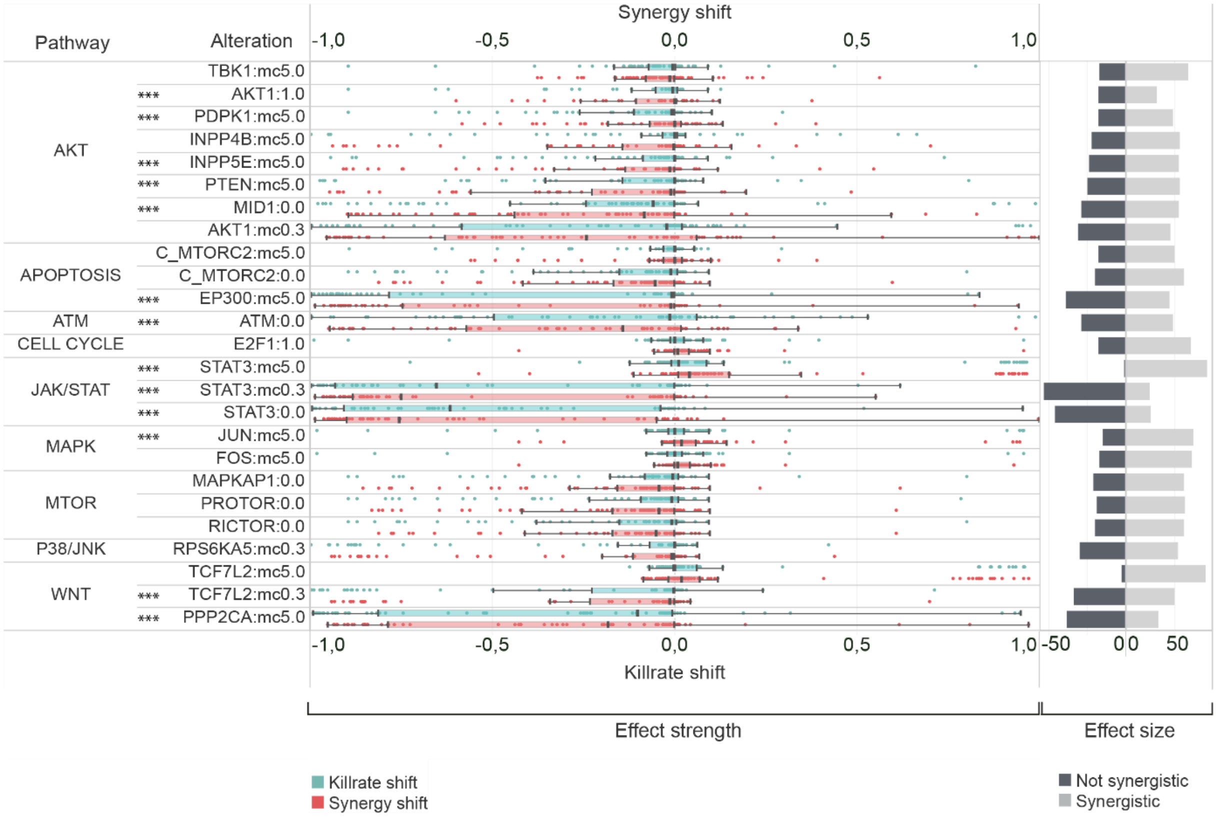

**Supplementary Text Fig. 2. Effect strength and effect size of combination specific biomarkers for the PRKDCi:NFKBi combination.**

Figure represents the statistically significant ($p<0.001$) shift of killrate and synergy influencing biomarkers in cell lines. Horizontal bars demonstrate the sample size for each biomarkers, while the distribution of effect strength components (killrate shift and synergy shift) are represented in boxplots. Abbreviation: *** p≤0,001 (rest of the visualized biomarkers statistical significance's is p≤0,05, 0.0 – loss-of-function, 1.0 – gain-of-function, mc5.0 – overexpression, mc0.3- underexpression. "C" prefixes represent protein complexes in the network.

#### Discussion of combination specific biomarkers

We observed that this combination has an indication specific synergy pattern. Unfortunately, as a novel combination prediction there were no experimental synergy scores available for comparison, and the literature evidence is also very limited. PRKDC (or DNA-PK) is a DNA-dependent protein kinase with a key role in repairing double-stranded DNA breaks, caused by DNA-damaging agents through non-homologous end joining (NHEJ). PRKDC is constituted from catalytic subunits (DNA-PKcs) and Ku70/80 DNA-binding subunits (Chang et al., 2017), and phosphorylates proteins like transcription factors, RNA polymerases, p53 or Ku70/Ku80 (XRCC6/5) (Mohiuddin and Kang, 2019).

Studies reported that in inflammatory diseases PRKDC is required for the expression of nuclear factor κB (NFKB) target genes that are downstream of TNF-α (Ghonim et al., 2015). Since inflammation is an important hallmark of especially metastatic cancer progression (Colotta et al., 2009; Grivennikov et al., 2010), the interplay between PRKDC, NFKB, TNF-α and interleukins could be a relevant approach for cancer cells in modifying their external microenvironment to increase the possibility of cell survival (Colotta et al., 2009; Yu et al., 2007). Therefore, understanding molecular alterations affecting synergy could be a viable approach to identify sensitive cell populations to this combination.

Based on our biomarker screen STAT3 and JUN overexpression are potentially relevant biomarkers of combination sensitivity. The role of STAT3 and JUN overactivation in connection with the NFKB signaling is controversial in the literature, described either as an oncogenic or as a tumor suppressor, leading to a context dependent effect on cell survival (Chaudhary et al., 1999; Huynh et al., 2019). In our model overexpression of STAT3 and JUN significantly increased synergy and increased the cell killing effect of this combination in some cells. On the contrary, both the inactivation and under-expression of STAT3 caused the opposite effect in the majority of the cell lines, leading to decreased synergy and cell killing. Therefore, based on our modeling STAT3 is a strong hypothesis in influencing synergy between these two compounds despite the fact that the effect of STAT signaling is controversial. On one hand, it has the ability to stimulate cell proliferation by activating CCND1. On the other hand, it can elevate the risk of aneuploidy, which can be the trigger point for the cell to start its mechanism of delayed mitosis-linked cell death. STAT3 can also activate anti-apoptotic proteins as well, like MCL1, BCL2, BIRC5. Thus, whichever scenario is about to happen it depends on the downstream members of STAT signaling and not on STAT3 itself.

Similar effect was observed after the inhibitory perturbation of NHEJ cross-talk pathway members, such as ATM or PPP2CA and TCF7L2 from the WNT pathway, resulted in weaker combination synergy. PPP2CA is an element of the PPP-family phosphatases. They regulate proteins by dephosphorylation included in cell cycle, DDR and several kinetochore kinases (Janssens and

Goris, 2001; Watkins et al., 2012). TCF7L2 modification is known to have a negative effect on cell proliferation through the regulation of WNT. Disruption of beta-catenin/TCF7L2 assembly's activity in colorectal carcinoma cells induces a rapid G1 arrest and blocks a genetic program that is physiologically active in the proliferative compartment of colon crypts, which is consistent with our results, showing that TCF7L2 under-expression has cell survival decreasing effect, beside its synergy lowering power (van de Wetering et al., 2002).

### Supplementary Methods

#### Supplementary Methods S1: Cell line and drug properties

##### Cell line datasets

97 cell lines were derived mainly from tumors of the breast (N = 27), lung (N = 27), bladder (N = 12) skin (N=9), and some other both primary and metastatic sites (N=22). Information about cell line characteristics were collected from the following databases: Expasy – Cellosaurus database (SIB Swiss Institute of Bioinformatics), ATCC, DSMZ (Leibniz Institute) (Supplementary Data S1).

Screening for synergistic combination identification included 58 compounds covering broad spectrum of mechanism of action groups, from which 12 was acting on DDR as main drugs. Out of the other paired combination drugs 18 targeted receptor and non-receptor tyrosine kinases (TK/RTKs), 14 targeted some elements of the PI3K-AKT-mTOR signaling, and 4 was for inhibiting apoptosis and ERK. The remaining drugs targeted pathways, such as JAK-STAT, or NFKB (Supplementary Data S1).

##### Compound databases

Benchmarking our *in silico* predictions require reliable and commercially accessible sources of pharmacokinetic and pharmacodynamic data. Therefore, we collected information on compound target profiles using ChEMBL and cellular response for compounds from databases such as Cancer Cell Line Encyclopedia, Cancer Therapeutics Response Portal, FIMM, GCSI, Genomics of Drug Sensitivity in Cancer, Oncolines, PharmacoDB and we also used the available DREAM *in vitro* data (Supplementary Data S1).

##### Determination of genetic variants

For the genetic variant annotations the Ensembl Variant Effect Predictor (VEP) was used with the parameters below, where input_file is a VCF format and the output_file is the standard output of the tool.
*Vep –af –af_1kg –af_esp –af_gnomad –appris –biotype –canonical –ccds –check_existing –distance 5000 –domains –hgvs –numbers –plugin dbNSFP, dbNSFP.gz, codon_degeneracy,*

*MetaSVM_score, MetaSVM_rankscore, MetaSVM_pred, MetaLR_score, MetaLR_rankscore, MetaLR_pred, Reliability_index –polyphen b –protein –pubmed –regulatory –sift b –species homo_sapiens –symbol –tsl –12niport –cache –input input_file –output output_file*

### Supplementary Methods S2: Description of *in silico* simulation types

#### Native simulation

To generate a cell population, consisted from 100 individuals cells from a single simulated cell, minor random perturbation are added to the mutational input layer of the simulation as a representation of both genotypic variance and therefore phenotypic heterogeneity inside the cell population. Perturbations are affecting the activity and concentration parameters of the nodes. Cell line specific phenotypic behavior was measured at the end of the simulation by aggregating the corresponding cell fate scores, such as cell cycle and cell death of the 100 attractors in the virtual population. We interpret our results as population based values. Thresholds for the combination of cell fate scores define the viability verdict of the cell population. If the cell population's viability score is above 0.6, we label it as alive. In case of dead cell line the same score is under 0.4. Marginal scores between these two cut off values representing cells with unknown behavior.

#### Monotherapy response screening and calculation of *in silico* IC50 values

The setup of cell line's to match their native behavior is followed by the calibration of their drug response to match their *in vitro* counterparts accurately. Modifying the parameters of a node and its input edge weight makes it less or more dependent on the upstream regulator node. Monotherapy response measurement is based on the ratio of alive and dead cells contained in the virtual cell population, therefore normalized survival and cell death scores are calculated. The overall survival (S) of cell lines and killrate of a given dose of drugs are calculated with the following equation:

$$S = \frac{relative\ cell\ cycle\ index}{1 - relative\ apoptosis\ index}$$

$$Killrate = 1 - S$$

We retrieve the final IC50 value by fitting a three-parameter Hill function on the results. The point on the dose axis where the fitted curve reaches a killrate of 0.5 is the *in silico* IC50 value for the tested drug on the tested cell line.

#### Combination therapy screen

In case of resistance or high toxicity for a specific drug, combination therapy is applied with at least two, potentially synergizing drugs. To find out which drugs would benefit patients the most and how they affect the *in silico* cells we created the combination screen where two sets of drugs are tested on the same cell line with either a symmetric or asymmetric dose grid. Every cell line - combination pair are tested in a full dose grid, and Bliss synergy score is calculated for all points of the grid.

##### Processing raw data of combination synergy

Let $K_a$ and $K_b$ denote the proportions of cancer cells that died following drug treatment a and b, respectively, where $\{K_a, K_b \in R \mid 0 \leq K_a, K_b \leq 1\}$. The model states that if drug a and b acts independently the predicted combination effect of the two drugs $\widehat{K}_{ab}$ can be calculated as:

$$\widehat{K}_{ab} = K_a + K_b - K_a K_b$$

Let $K_{ab}$ represent the observed combination effect of drug a and b where $\{K_{ab}: 0 \leq K_{ab} \leq 1\}$, then "excess over Bliss" namely Bliss score $BS$ can be written as:

$$BS = K_{ab} - \widehat{K}_{ab}, \{BS: -1 \leq BS \leq 1\}$$

where BS > 0 indicates synergy and BS < 0 indicates antagonism.

*Bliss max definition:*

*Bliss max* is the maximum over all of the Bliss scores in a combinational grid. Let

$$v = \{v_i \in R_{>0}, i = 1, \dots, n_a\}$$
$$u = \{u_i \in R_{>0}, i = 1, \dots, n_b\}$$

denote the sets of doses applied to the cell as combinations from drug a and b, then the Cartesian product $v \times u$ is the set of all ordered dose pairs from drug a and b. Let

$$bm = (bm_1, \ldots, bm_{n_a x n_b})$$

denote the vector of Bliss scores where each element $b_i \in bm$ is the Bliss score of the corresponding dose pair of $v \times u$, then "*Bliss max*" is the maximal value of vector bm:

$$Bliss\, max = max\ bm_i, i = 1, \ldots, n_a x n_b$$

Taken the Bliss scores in a subgrid of combinational grid where subgrid is determined by the monotherapy IC50 values of the two compounds *Bliss max ic50* is the maximum Bliss score over this subgrid. In a given combinational grid *Maxdose* determines which dose pair of drug A and B have the maximum Bliss score of the IC50 subgrid and *Bliss ic50 killrate* is the killrate of drug combination at *Maxdose*.

Let $IC50_a$ and $IC50_b$ represent the half maximal inhibitory concentration of drug a and drug b respectively and A denotes the following subset of $v \times u$:

$$A = \{(n, m): (n, m) \in v \times u, n < IC50_a, m < IC50_b\}$$

Let $bm_{ic50}$ denotes the vector of Bliss scores where each element $bm_{ic50}(i) \in bm_{ic50}$is the Bliss score of the corresponding dose pair of A, in this case "*Bliss max ic50*" is defined by

$$Bliss \max ic50 = max\ bm_{ic50}(i) \ \ i = 1, \ldots, n$$

and *Maxdose* is the corresponding ordered dose pair of Bliss max ic50 from set A and *Bliss ic50 killrate* is the observed killrate of drug combination at *Maxdose*.

##### Quality control of raw combination results

Cell line specific combination grids were gone through a quality control process in which we checked the combinational grids for non-monotone dose response issues focusing on the subgrid area. This area is determined by the monotherapy IC50 values of the two compounds. Combination grids where the normalized survival values were not decreased according to increasing drug doses were judged individually and distinguished into the following categories:

(i) Where normalized survival values varied by a minimum of 0,2 in case of both compounds specifically on the subgrid area, the combination grid was considered to be not reliable and discarded from further analysis.

(ii) Combinations where the same phenomenon could be observed, but above the subgrid area, were labelled as uncertain.

(iii) In case of observing invers association between increasing drug doses and cell viability, the combinations were considered to be reliable.

We proceeded to further analysis only with combinations tagged as uncertain and reliable. To determine cut-off values to distinguish between synergistic and non-synergistic combinations, we applied a K-means clustering based algorithmic method. Similar approach has been already used in the literature to discretize gene expression data (Gallo et al., 2016) (Supplementary Fig. S4).

#### Biomarker screen

The monotherapy response screen can be supplemented with extrinsic, artificial modifications of node activity and concentration parameters. This simulation type examines the power of the added alterations on shifting of monotherapy IC50 values. Inactivating (loss-of function) mutations result in the possible decrease of maximum activity of a protein to zero. Activating mutations (gain-of-function) mutations constitutively activate a protein by increasing the possible minimum activity of a given protein to one. Protein abundances in the Simulated Cell are between 0 and 3, with most values being in the range of 0-1. An average 0.6 fold-change value reaches the potential maximum when multiplied by 5, while the same value goes to 0.2 (which does not completely turn the protein off, but already significantly impacts its ability to pass signals downstream) when multiplied by 0.3. Therefore, to generate overexpression or underexpression in the network, the actual concentration parameter of a given node is multiplied by 5 or 0.3 in every time-step, respectively. A monotherapy specific biomarker is defined as the alteration of a specific protein which results in the IC50 shift of a tested drug as monotherapy, whereas combination specific biomarker is an alteration that modify the synergy of a combination therapy but failed to cause shift in the monotherapy response.

##### Systematic prescreen for verdict changer biomarkers

The systematic prescreen was performed on 13 cell line models for AZ13150560:AZ12879988 combination and on 6 cell line models for Olaparib:AZ13535704 combination. Both sample set was chosen to represent both synergistic and not synergistic combination-cell line pairs, to make the detection of biomarkers possible with synergy increasing and decreasing character. The prescreen searched for verdict changer molecular alterations that shifted the treated simulated cell's viability verdict status from alive to dead or vice versa, for sensitivity or resistance biomarkers, respectively. The prescreen systematically simulated the effect of loss-of-function, gain-of-function, underexpression and overexpression of each node in the network while adding single compound or compound combination effect to the simulation on doses manually curated for all cell lines. The doses were defined to have the *in silico* cell's viability clearly in the alive or dead verdict and to be in the most synergistic dose-pair if that coupled with decreased viability, the general concept during manual dose selection is to define doses that are 3-times higher or lower than the *in silico* cell lines IC50s.

Combination specific verdict changer alterations were selected if they shifted the viability on the specific cell lines only by adding the compound combination and did not shift the viability under monotherapy. For the AZ13150560:AZ12879988 combination we further narrowed down the combination specific verdict changer alterations by selecting those for further detailed screen that are members of mTOR (TOR signaling), AKT (3-phosphoinositide-dependent protein kinase activity), JAK-STAT (receptor signaling pathway via JAK-STAT), PLCG (phospholipase C activity), TGFB (transforming growth factor beta receptor signaling pathway), TNF (tumor necrosis factor-mediated signaling pathway), p38-JNK (stress-activated MAPK cascade), NHEJ (double-strand break repair via nonhomologous end joining), NFκB (I-kappaB kinase/NF-kappaB signaling) and apoptosis pathways based on our network annotation. These steps resulted in 41 alterations (8 sensitivity and 33 resistance biomarker candidates) for Olaparib:AZ13535704 and

166 alterations for AZ13150560:AZ12879988 (120 sensitivity and 46 resistance biomarker candidates) to proceed with for the detailed screening.

#### Screen for combination specific synergy shifting biomarkers

In order to identify synergy-related biomarkers we compared the combined drug efficacy with and without added extrinsic mutations (henceforth *nomut* and *mut*, respectively). Two of the main metrics was *Bliss max ic50 nomut* and *Bliss max ic50 mut* which are the Bliss max ic50 values of the unmutated and mutated counterpart of a combination drug experiment. Comparison of these is an indicator of the mutation effect on the synergistic behavior of the drug combination.

Beside the specific synergy metrics we also examined the killrates of the combination drugs, namely *Bliss ic50 killrate nomut* and *Bliss ic50 killrate mut* which are the *Bliss ic50 killrate* of the unmutated and mutated counterpart of drug combination.

### Supplementary Methods S3: Analysis of features governing *in silico* predictivity

The goal of the analysis was to create a model which captures the association between the characteristics of cell lines and drugs and the predicted synergy by our network. The used model should be interpretable hence the aim of this exploration was "inference" focused. The main idea was to use features of cell lines/drugs which were used as input data layer for the Simulated Cell network.

#### Identification of features important to predict synergistic combinations

**The used datasets included**

- Output of the postprocessed combination results, which aggregates the dose grids into single, relevant metrics (Supplementary Data S2)
- Binarized Bliss_max_IC50 values were used as a measure of synergy.
  Thresholds: > 0.25 for synergistic, < 0.25 non-synergistic (Supplementary Fig. S4)
- 59947 unique cell line-combination drug pair described with max, average Bliss scores and other features (Supplementary Data S4)
- 97 cell lines, 684 drug combinations (Supplementary Data S1)

**Input features**

1. Monotherapy related features
    a. Modelling feasibility (Supplementary Data S1)
    b. Fold accuracy (considering AZ in vitro results as gold standard).
    When there were no cell line-drug match with any DREAM results we replaced the missing values with the average IC50 value of the drug in all other cell lines.
    c. Target number in the signaling network and not in the signaling network, with various threshold values (100, 1000, 10000 nmols)
    d. Two kind of mechanism of action (MoA) categories (Supplementary Data S1)
    e. Monotherapy cell line coverage of the different drugs (Supplementary Data S1)
    f. Number of common targets in the signaling network
    g. Number of common targets in total

2. Compound target related features
    a. Median binding affinity values of proteins included in the signaling network
3. Mutation readiness
    a. All unique included proteins with their corresponding encoded value (0: no mutation at this gene, 1: gain-of-function mutation, 2: loss-of-function mutation). (Described in Methods)
4. Fold change data
    a. Fold change value of the genes in the network (Described in Methods)

**Target feature**

- Binarized Bliss_max_IC50 values were used as a measure of synergy.
  Thresholds: > 0.25 for synergistic, < 0.25 non-synergistic (Supplementary Fig. S4)
  Categorical transformation was needed due to the result of comparison with DREAM's *in vitro* results.

**The created dataset included:**

- 20 monotherapy related features
- 262 compound target features
- 1681 fold change features
- 534 mutation readiness features

As we have high-dimensional input data features selection methods were used to reduce the dimensionality of the dataset and an embedded method was applied which both executes feature selection and model fitting. Logistic regression was used with elastic net-based penalty to shrink coefficients of irrelevant features towards zero. In order to evaluate the model properly we implemented several cross-validation scheme described by Preuer et al. (Preuer et al., 2018).

Evaluation on new drug combinations, new drugs or new cell lines which the model hasn't seen before shows much more realistic measure of the generalization capabilities of the model than random splits. In order to properly evaluate the generalization capabilities of the model we used the "leave drug combination out" cross validation method as Preuer et al. Hyperparameter optimization was used to choose the best performing elastic-net mixing parameter and inverse regularization strength parameter (Supplementary Methods S2 Table 1).

| **Parameter set (C, I1 ratio)** | **Average weighted F-1 score** |
|---|---|
| (0.001, 0.1) | 0.7955 |
| (0.001, 0.5) | 0.7342 |
| (0.001, 1.0) | 0.6825 |
| (0.01, 0.1) | 0.8353 |
| **(0.01, 0.5)** | **0.8368** |
| (0.01, 1.0) | 0.8155 |
| (0.1, 0.1) | 0.8174 |
| (0.1, 0.5) | 0.8206 |
| (0.1, 1.0) | 0.8100 |
| (1, 0.1) | 0.7860 |
| (1, 0.5) | 0.7884 |
| (1, 1.0) | 0.7758 |

**Supplementary Methods S2 Table 1.**
**10-fold ”leave drug combinations out” cross validation result of different hyperparameters.**

The first column contains hyperparameters (l1 ratio=elastic net mixing parameter, C= inverse of regularization strength) of the elastic net model, second column shows the average weighted F-1 scores of the fitted model with the given hyperparameters using the above described evaluation method.

Henceforth the best former parameter setting were used based on the maximum averaged F-1 score reached:

- 0.5 as elastic-net mixing parameter
- 0.01 as inverse regularization strength which mean "half-way between lasso and ridge penalty" and quite strong regularization

We standardized the features because introducing regularization term make it a scale-variant model. According to Supplementary Methods S2 Table 2 the model performs well on unseen drug combinations. Using the "leave combinations out" cross validation scheme we get a more realistic picture of the model coefficients. Inspecting the coefficients we can identify the relevant features and their impact.

| *10 fold "new drug combination" cross validation* | **Train mean** | **Train STD** | **Test mean** | **Test STD** |
|---|---|---|---|---|
| **Balanced accuracy** | **83,1%** | **0.4%** | **77.35%** | **1.96%** |
| **Matthews correlation coefficient** | **69.24%** | **0.6%** | **59%** | **3.2%** |

**Supplementary Methods S2 Table 2.**
**Classification result of logistic regression model with elastic net.**

The table shows the evaluation of the best performing model using the "leave combinations out" cross validation scheme measured by various classification metrics (balanced accuracy, Matthews correlation coefficient).

#### Identification of differences between correctly and wrongly predicted synergies based on monotherapeutic features

**Input features:**

- Modelling feasibility (Supplementary Data S1)
- Fold accuracy (considering DREAM in vitro results as gold standard) (Supplementary Data S1)
- Number of targets included and not included in the signaling network with various threshold values (100, 1000, 10000 nmol)
- Two kind of *in silico* mechanism of action (MoA) categories (Supplementary Data S1)
- Cell line coverage of the different drugs (Supplementary Data S1)
- Number of common targets included in the signaling network
- Number of common targets in total

**Target feature:**

- Correctness of our prediction (compared to the DREAM in vitro synergy data)

  Correctness was quantified based on thresholded DREAM *in vitro* synergy scores and our Bliss IC50 maxes (IV threshold: 30, IS threshold: 30) (Fig. 4B)
- Using these thresholds we got around 66 % balanced accuracy and the below confusion matrix (Supplementary Methods S2 Table 3)

| Labels | Predicted Synergistic | Predicted Non-Synergistic |
|---|---|---|
| Actual Synergistic | TP = 56 | FN = 72 |
| Actual Non-Synergistic | FP = 138 | TN = 711 |

**Supplementary Methods S2 Table 3.**
**Confusion matrix of predictions taking DREAM results as true labels.**

TP – True positive, TN-True negative, FP – False positive, FN – False negative

Since we cannot observe any input feature correlating with our target feature nor any meaningful differences between the classes of input features by visual inspection, we fitted linear and non-linear models to classify our prediction categories. We experimented with all confusion matrix elements (TP, TN, FP, FN) and binarized categories as well (Correctly-Not correctly predicted). Even with handling class imbalances these models were not satisfying as TN samples over-dominated all classifiers. The exclusion of TN samples from the data caused significant performance improvement (Supplementary Methods S2 Table 4). These TN samples couldn't be separated by using our independent variables.

| *10-fold cross validation scores with RF* | **Mean balanced accuracy** | **Std of balanced accuracy** |
|---|---|---|
| **Inclusion of TN samples** | 55.37 % | 8.3 % |
| **Exclusion of TN samples** | 66.09 % | 3.53 % |

**Supplementary Methods S2 Table 4.**
**Balanced accuracy including-excluding TN samples.**

Mean and standard deviation of balanced accuracies evaluated using 10-fold cross validation with vs without true negative samples.

To identify only the relevant features we applied Recursive Feature Elimination to discard irrelevant input features. As we can inspect, including other features beside the 4 most relevant feature doesn't really increase our performance metrics (Supplementary Fig. S9). Therefore, we continued modelling with excluding the least important features (feasibility and MoA category labels) and used the top 9 features and hyperparameter optimized a Random Forest model (Supplementary Methods S2 Table 5).

| *10-fold cross validation scores with RF* | **Mean balanced accuracy** | **Std of balanced accuracy** | **Mean weighted F-1 score** | **Std of weighted F-1 score** |
|---|---|---|---|---|
| **Without hyperparam opt** | 69.51 % | 7.872 % | 68.92 % | 5.01 % |
| **With hyperparam opt** | 73.47 % | 13.56 % | 74.25 % | 11.64 % |
| **With optimized hyperparam and oversampling** | 76.68 % | 4.8 % | 76.46 % | 4.55 % |

**Supplementary Methods S2 Table 5.**
**Hyperparameter optimization and oversampling results.**

The columns denote the various classification metrics used for evaluation, while the rows are models with different conditions.

According to the gini index based feature importance values of the optimized Random Forest (RF) model, fold accuracies, cell line coverage and the number of common targets seems to be important as well for the model (Supplementary Fig. S7 and Supplementary Methods Table 6).

| **TOP relevant features in decreasing order** |
|---|
| Drug A fold accuracy |
| Drug B fold accuracy |
| Cell line coverage |
| Drug B MoA label |

**Supplementary Methods S2 Table 6.**

**The most relevant features based on RFE.**

The table includes the 4 most relevant features in decreasing order based on Recursive Feature Elimination with random forest model evaluated by cross-validation. Other features were eliminated due to their marginal/non-existent contributions to model performance.

In order to further investigate how these features impact our target feature, the correctness of our predictions, the game theory based SHapley Additive exPlanations (Lundberg, 2017) was used. In order to simplify the interpretation of fold accuracy variables we took the absolute value of them. As we can inspect (Supplementary F8), low value of *cell_line_coverage* increase the probability of wrong prediction category, while high value of *cell_line_coverage* decrease the probability of correct prediction category. Similarly, low value of *drug_b_fold_accuracy* increase the probability of wrong prediction category, while low value of *drug_b_fold_accuracy* increase the probability of correct prediction category.

### Supplementary Figures

**Supplementary Fig. S1: Comparison of *in vitro* dose responses from different sources across the DREAM challenge cell line set**

Setting up the correct drug response within each cell line and drug pair is one of the most challenging but crucial part to prepare the in silico Simulated Cell for applying adequate prediction methods. Since most database contain one datapoint for each measurement it is hard to draw conclusion from mismatching in silico and in vitro responses. Hence, a tradeoff had to be made between the correct drug response prediction based on single datapoint and accurate underlying biology, especially where protocols, such as treatment times differ. The plot demonstrates nicely the differences between in vitro IC50 data from various external sources (showed in red color) and DREAM in vitro data where it was available (showed in turquoise color).

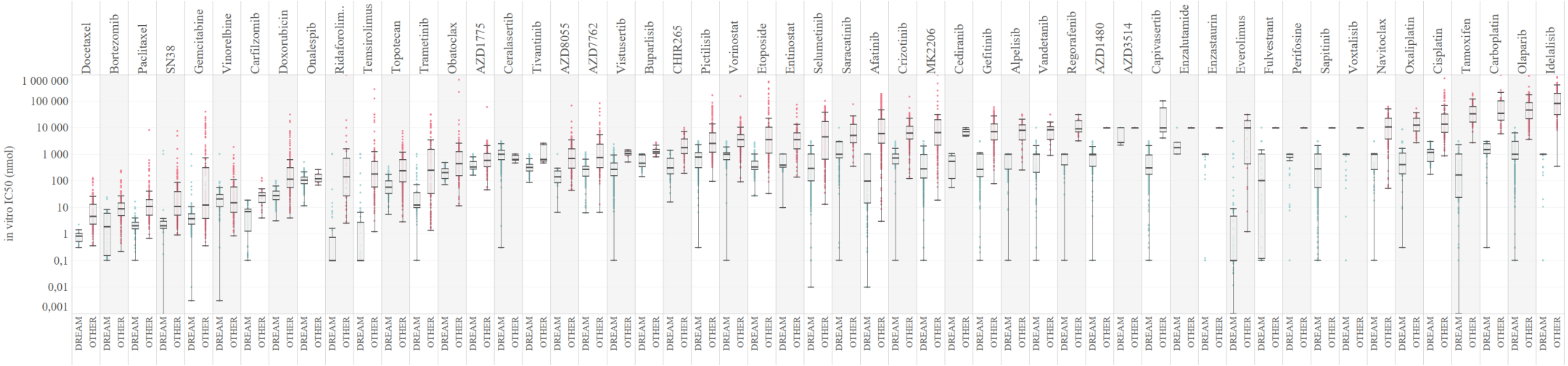

**Supplementary Fig. S2: ROC curves of monotherapy simulations for compounds targeting DDR or non-DDR MoA categories**

Results were aggregated based on the targeted MoA category of compounds, resulting in DDR and non-DDR groups. AUC metric is indicated.

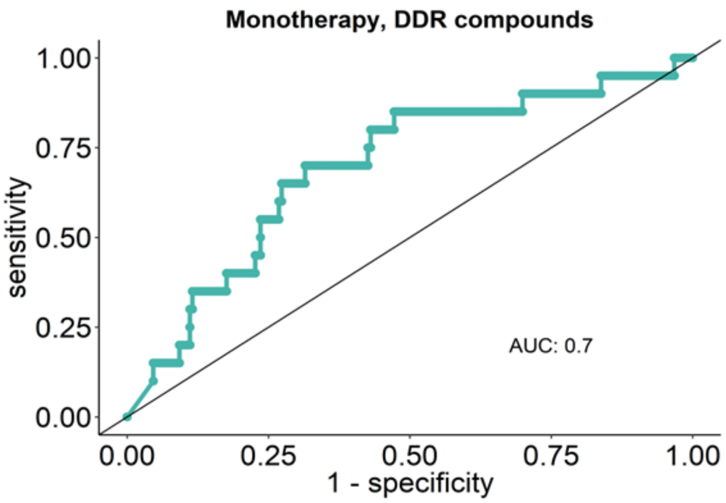

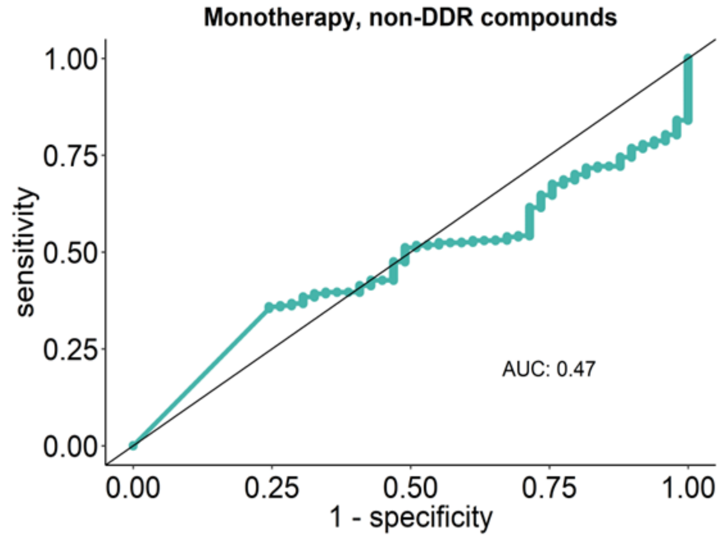

**Supplementary Fig. S3: Examples of different combination grid quality categories**

Each dose grid represents normalized survival values in each dose points. A) Unreliable combination: possible false positive synergy scores due to non-monotone dose response with both compounds under their IC50. 0.17% of all combination were considered to fit into this category. B) Uncertain combination: possible false positive synergy scores due to non-monotone dose response with both compounds over their IC50. 5.43 % of all combination were distinguished as uncertain combination. C) Reliable combination: synergy score is considered reliable based on the preliminary quality check of the survival score landscape on the specific cell-drug combination dose grids. 65.96% of the combination were labelled as reliable based on the combination grids. The remaining 28.43% were considered to be not-synergistic, since one of the combination partner's effect dominated the observed effect, the other partner did not have the space for showing an effect on survival, example not shown. Abbreviations: AZ_AKT: AKT inhibitor, AZ_PRKDC_I: PRKDC inhibitor 1, AZ_PRKDC_II: PRKDC inhibitor 2, AZ_PI3K- Protein synthesis (PI3K) inhibitor (molecular target profiles were provided by AstraZeneca).

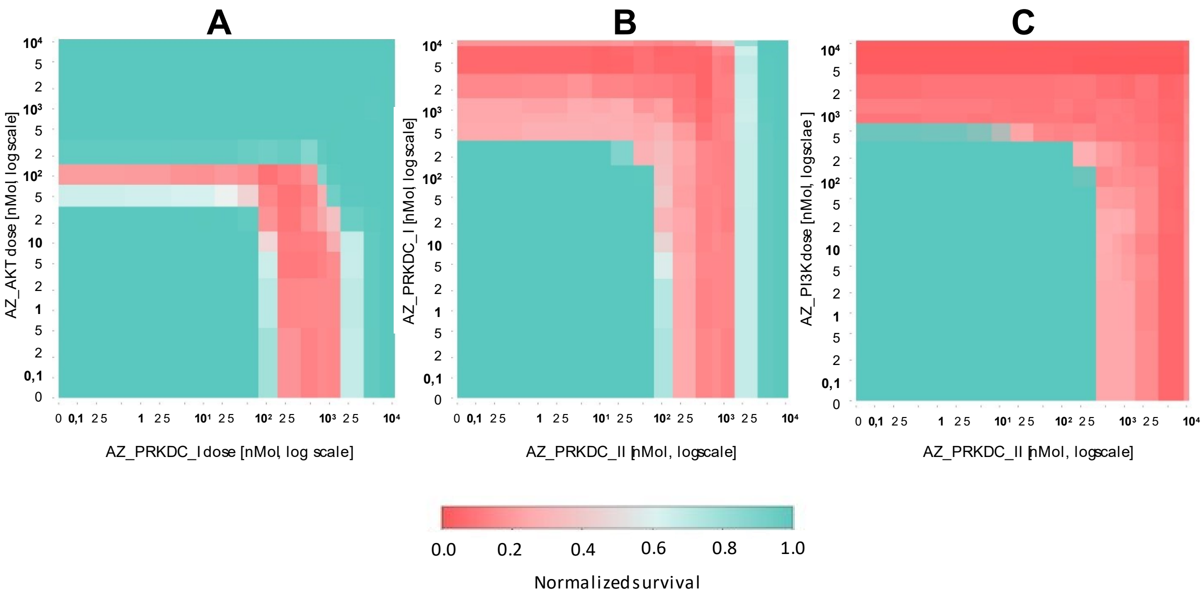

**Supplementary Fig. S4: Discretization of synergy scores with K-means clustering**

An algorithmic method was applied to discretize the continuous synergy scores. Thresholds for different synergistic categories were established using k-means clustering, a widely used discretization method in biological data analysis. Our assumption is that the distribution of synergy scores is the mixture of different synergy categories distributions. After the evaluation of the clustering results we draw two thresholds splitting the combination-cell line pairs into three distinct categories such as non-synergistic (Cluster 0), moderately synergistic (Cluster 2) and strongly synergistic (Cluster 1). For further evaluation we prioritized the most promising, strongly synergistic pairs from Cluster 1.

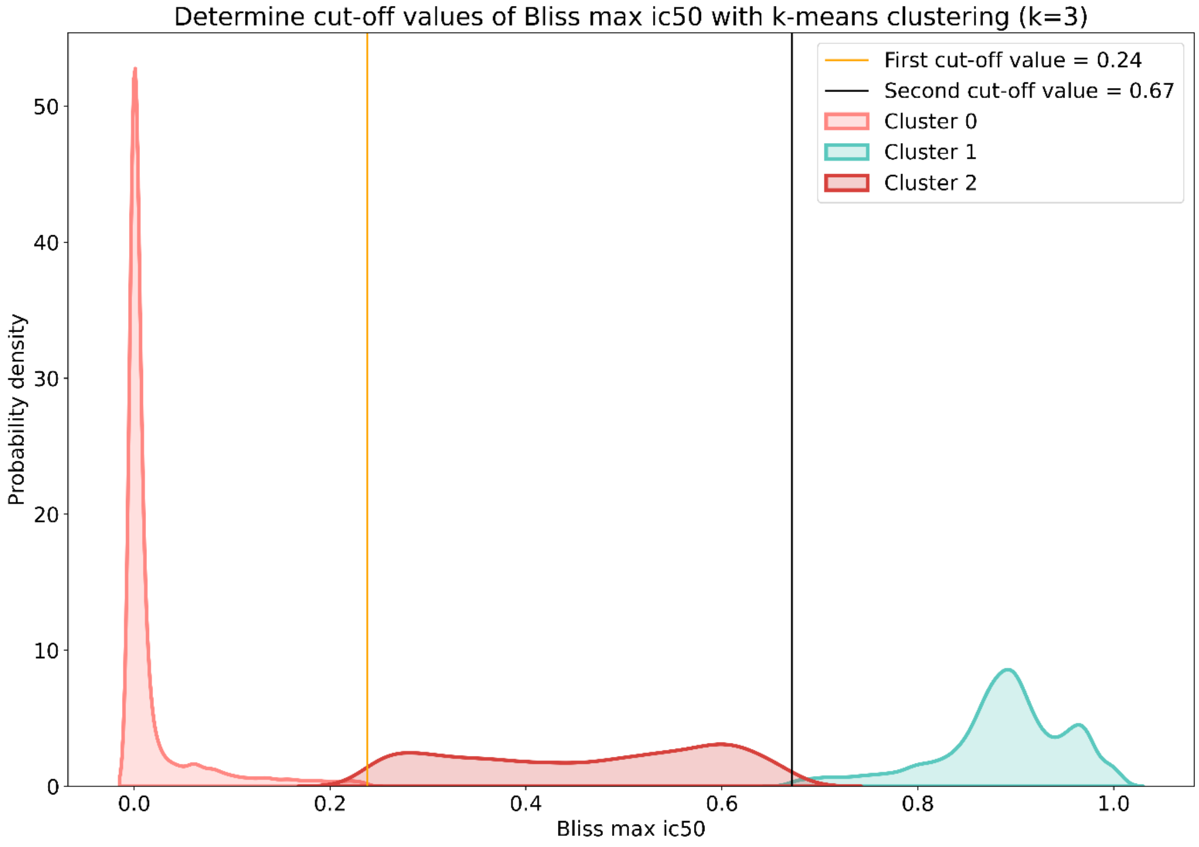

**Supplementary Fig. S5: Grouped indication specific landscape of DDR:DDR combinations on combination modified killrates and overall synergy**

The radar charts are representing different synergistic and non-synergistic combination pairs. Synergy is concluded from the average Bliss score with additional average killrate values. Higher the synergy is, the outer radius of the circle is reached at given indication group. Mean synergy and mean killrate were calculated for each MoA groups for all cell lines categorized into indication groups.

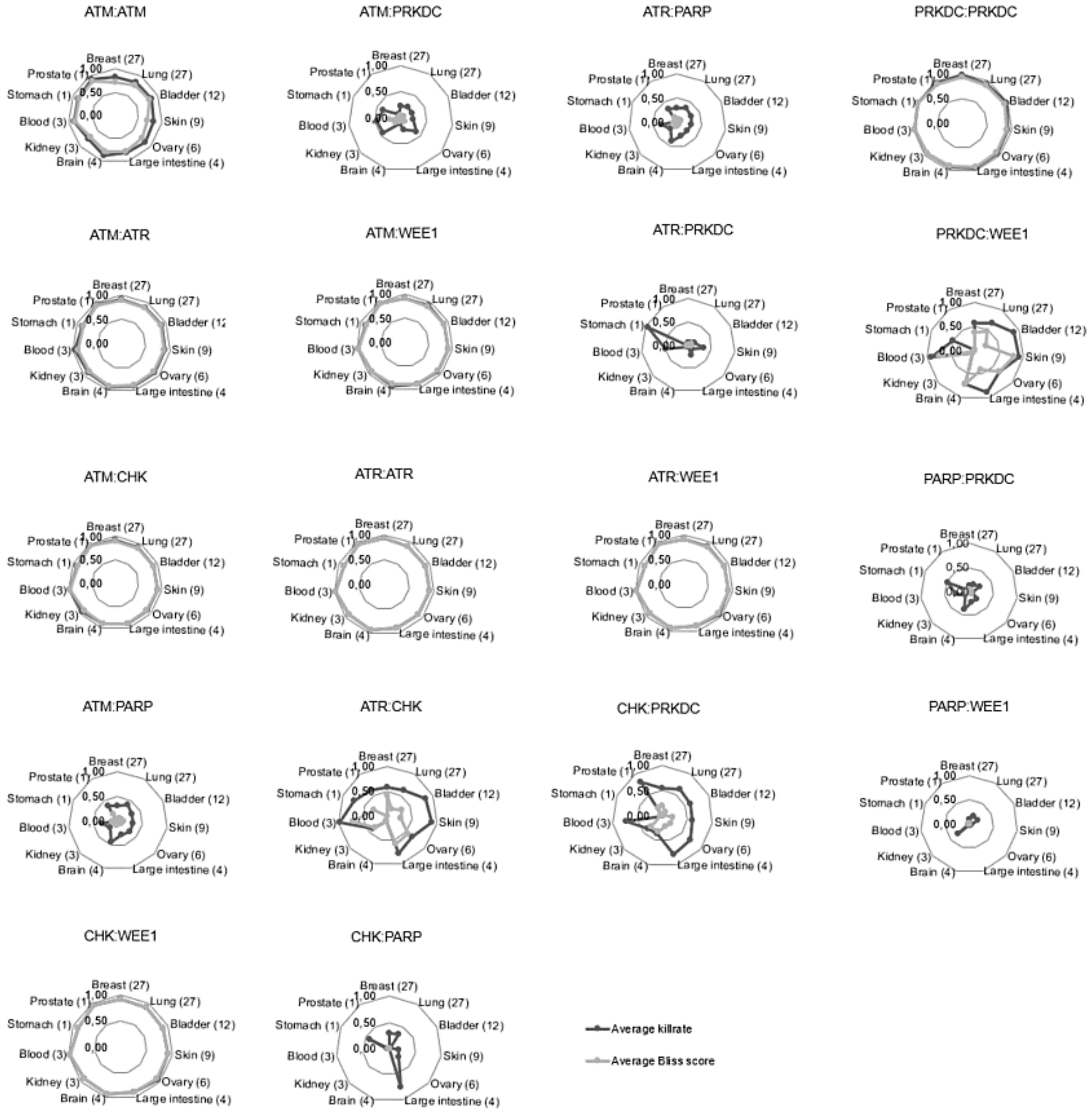

**Supplementary Fig. S6: Performance analysis of combination synergy benchmark**

A) Overall performance analysis of combination benchmark consisting of 977 overlapping datapoints with the DREAM challenge in vitro synergy scores. For the AUC evaluation in vitro (IV) threshold 30 and in silico threshold 20 were used. AUC metric and the amount of synergistic simulations (syn) are indicated.

A

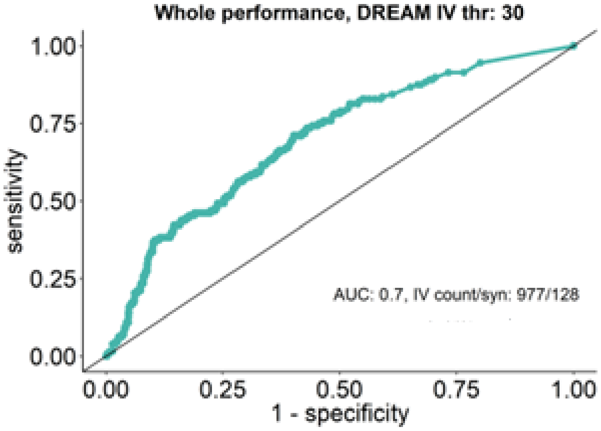

B) AUROC scores for the prioritized compounds based on the combination benchmark. The number of involved datapoints are indicated in parentheses. The in vitro threshold of 30 and in silico threshold of 20 were applied in the performance.

B

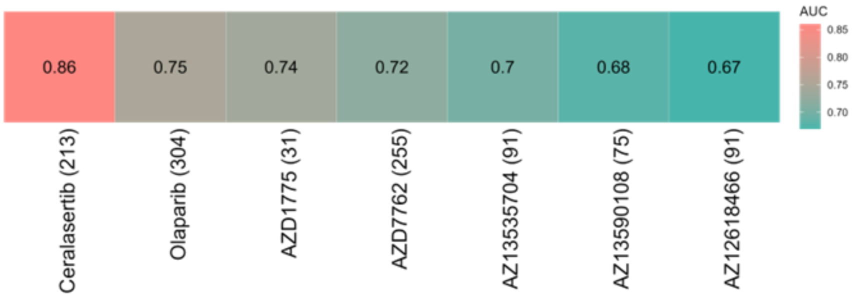

**Supplementary Fig. S7: Tree-based feature importance of association between monotherapy related features and combination prediction accuracy with a Random Forest model**

The examined features were: cell_line_coverage: coverage for the in vitro synergy of the combinations, drug_b_fold_acc: abs(log10(in silico IC50/AZ in vitro IC50) for the second compound (DDRi or nonDDRi) of the combination, no_of_common_targets_insig: number of common targets of drug_a and drug_b in the signaling network, no_of_common_targets: number of common targets of drug_a and drug_b, not only in the signaling network, drug_a_fold_acc: abs(log10(in silico IC50/AZ in vitro IC50) for the first compound (DDRi) of the combination, drug_a_target_num_ratio and drug_b_target_num_ratio: ratio of the modelled targets, drug_a_MoA_labels: MoA categories of the DDRi-s.

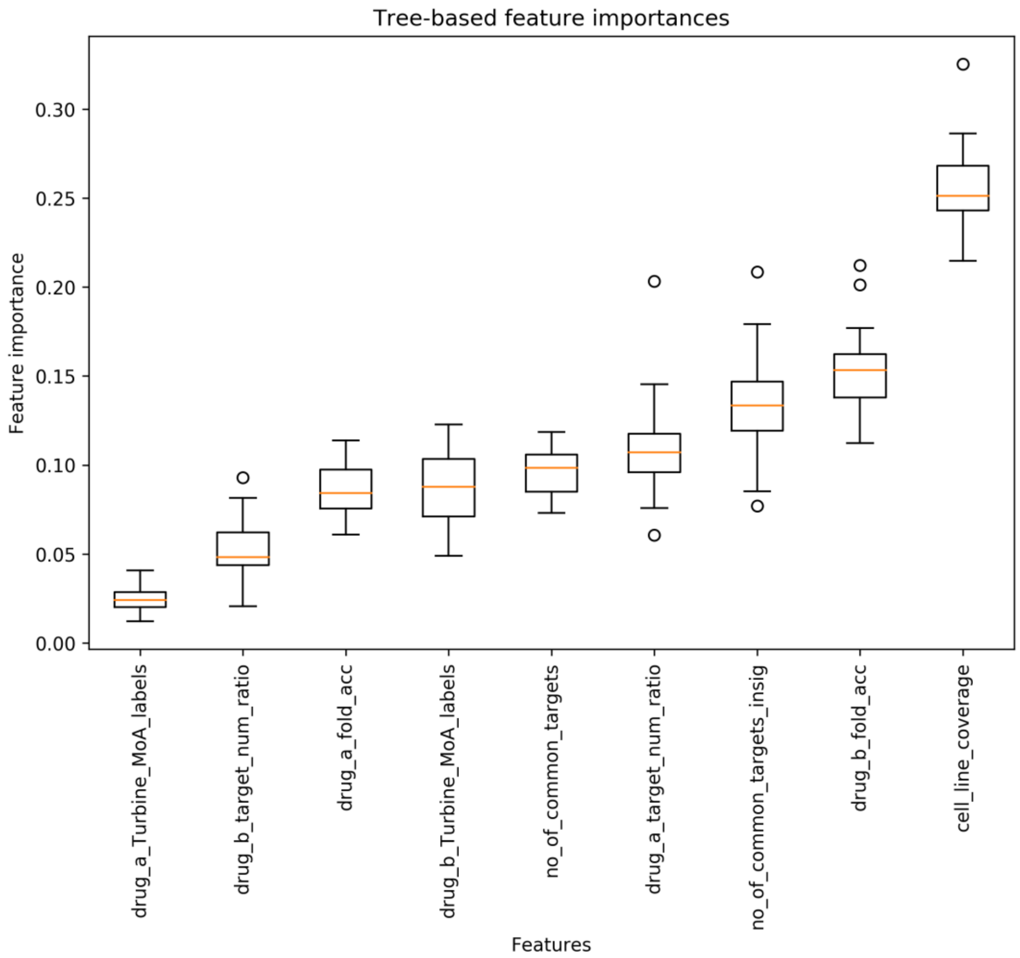

**Supplementary Fig. S8: Investigation of relevant features impacting synergy as a target feature using SHapley Additive exPlanations (SHAP)**

SHAP values of A) wrong predictions (FP, FN) and B) correct synergy predictions (TP). Game theory based SHapley Additive exPlanations (Lundberg, 2017) was used to further investigate the relationship of the features and the model. We can plot the SHAP values of every feature for every sample. The plot sorts features by the sum of SHAP value magnitudes over all samples, and uses SHAP values to show the distribution of the impacts each feature has on the model output. The features used here were "fold accuracy" for each drugs that indicates the magnitude of difference between in silico and in vitro data, "MoA labels" for each drugs which annotates the mechanism of each drugs into Mechanism of Action groups, the "cell line coverage" which counts the number of cell lines for one combination therapy we had data for, "no common targets" and "no common targets insig" counts the number of targets shared between the two drugs of a given combination all together and with those targets that are present in our network only, respectively, and finally the "target num ratio" as the percentage of given drug's target proteins are present in our network in a dose dependent manner, where off-targets are counted as well. The color indicates the feature value with red for higher and blue for lower values.

A

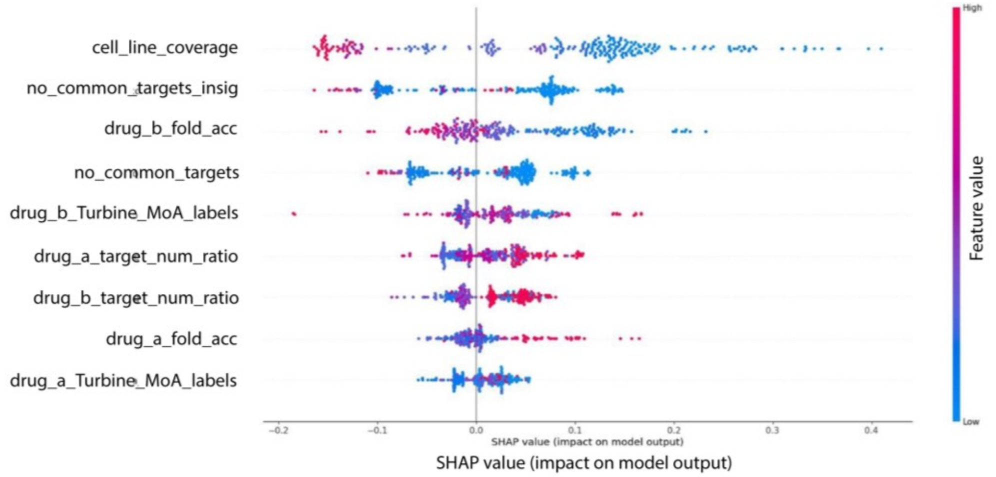

B

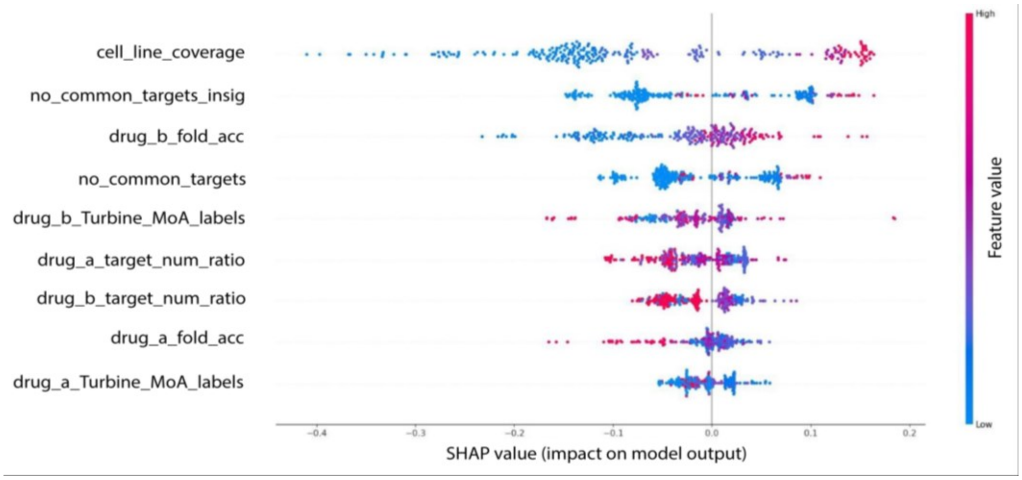

**Supplementary Fig. S9: Determining features of predicting combination synergy and their positive and negative coefficients from logistic regression model based on 10-fold leave drug combinations out cross validation**

The plots below show the top 20-20 variables with the largest and lowest odds ratios, respectively, to indicate the most relevant features and their relationship with the target in the logistic regression model.

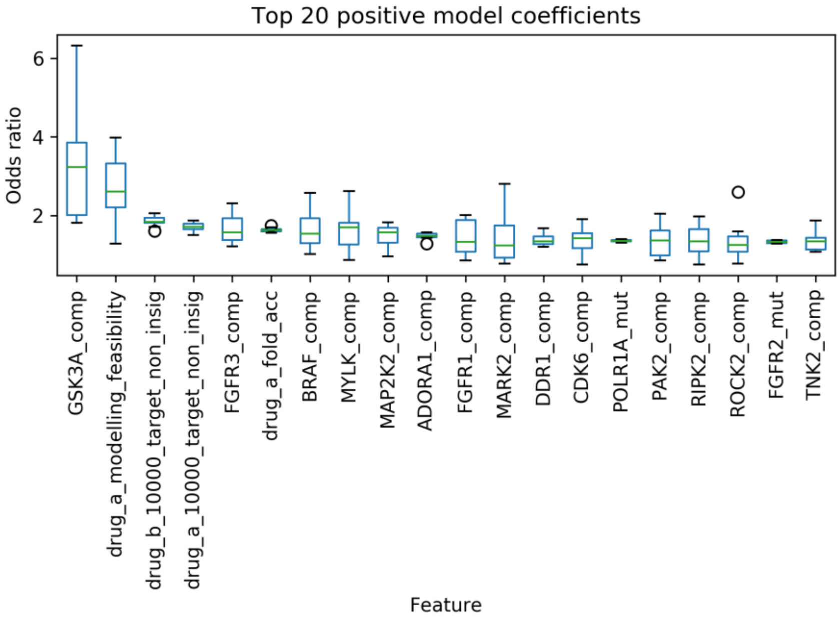

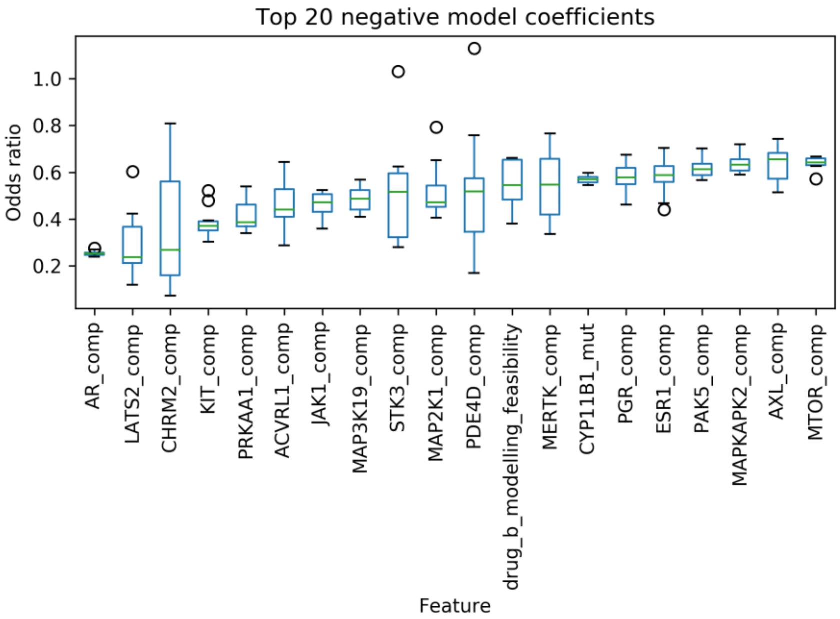

### Supplementary Data

#### Supplementary Data S1: *In silico* vs *in vitro* monotherapy measurements with respective cell line and compound annotation

This supplementary file contains the *in silico* IC50 prediction in nmol as in "IS (nmol)" coloumn for each drug – cell line pairs, annotated in "ID", "Cell line" and "Drug" coloumns. To have an experimental checkpoint for each prediciton we used the DREAM data where it was accessible or other open sources of *in vitro* measurements in nmol, stored in "IV DREAM (nmol)" and "IV OTHER (nmol)" coloumns, respectively. We also listed the one-fold accurate and inaccurate *in silico* predictions compared to their *in vitro* counterpart for both DREAM only and "Other" miscellaneous sources. Other databases are described in Supplementary Methods S3. To avoid cell line characterization issues, each cell line used in our screens are annotated with their metastatic status as in "Cell line origin", the source organ as in "Organ of origin", refered disease, MSI status and the adequate *in vitro* maintaining setting as in "Growth properties". For the compound annotation we found important to clarify the putative target and their respective targeted mechanism of action group, the feasibility score based on our internal benchmarking system and the cell line coverage we have data for for each drug listed.

#### Supplementary Data S2: Combination measurement metrics

This supplementary file contains each drug combination with all cell lines screened, i) the *in silico* IC50 value of each combination drug as monotherapy in given cell line (with values over 10,000 entered as 10,000), ii) the killrate value when the combination therapy was applied in given cell line at maximum Bliss synergy dose, iii) the maximum Bliss synergy score of the combination in given cell line and iv) the same with both compounds under the IC50 dose, and v) classification determining reliability of the combination's effect in given cell line.

#### Supplementary Data S3: Monotherapy biomarkers

To distinguish those biomarkers, which could modify the effect of the each drug alone from the combination specific ones, we screened for monotherapy biomarkers. Supplementary Data S3 contains every potential biomarker we predicted for each drug individually. Affected cell lines and their source of origin are stored as in "Cell line" and "Organ of origin ". Our predictions are coded as the gene and the effect of the mutation on given protein being loss-of-function (genename:0.0), gain-of-function (genename:1.0), underexpression (genename:mc0.3) or overexpression (genename:mc5.0) as in "Hypothesis" and "Hypothesis alteration". To ease the matching to different outer sources we listed the UniProt ID's of affected proteins as in "Hypothesis uniprot" and for easier search we also listed the gene name only from the hypotohesis as in "Hypothesis genename".

#### Supplementary Data S4: Combination biomarkers for PARPi:ATMi along with PRKDCi:NFKBi

Contains what we predicted as unique biomarkers for the combination therapy of PARP and ATM inhibitors and the combination of PRKDC and NFKB inhibitors for given cell line. Our predictions are coded as the gene and the effect of the mutation on a given protein being loss-of-function (genename:0.0), gain-of-function (genename:1.0), underexpression (genename:mc0.3) or overexpression (genename:mc5.0) as in “Hypothesis” and for easier search we also listed the gene name only from the hypotohesis as in “Hypothesis genename”. This data file also contains i) the killrate of the Simulated Cell without any treatment and with or without added alteration, ii) the maximum Bliss synergy score of the combination in the given cell line and the same with both compounds under the IC50 dose, iii) the *in silico* IC50 value of each combination drug as monotherapy in given cell line (with values over 10,000 entered as 10,000) and either with or without added alteration, iv) the magnitude of shift in IC50 for either drug by the hypothesis, v) sum of the Bliss difference matrix, which is the result of element-wise subtraction of non-altered cell line’s Bliss matrix from the altered cell line’s Bliss matrix, vi) sum of the killrate difference matrix, which is the result of element-wise subtraction of non-altered cell line’s killrate matrix from the altered cell line’s killrate matrix, and vii) the type of biomarker (sensitivity / resistance biomarker) based on overall_killrate_change.

#### Supplementary Data S5: The estimated patient population size for Olaparib:ATMi combination and the prevalence of the significantly strong synergy shifter biomarkers specific for the combination

To provide translatable combination specific biomarkers, we estimated the size of potential patient population who could benefit from the Olaparib:ATMi combination by calculating the frequency of patients with damaging BRCA1 or BRCA2 and ATM mutations, since these genetic factors are one of the main inclusion criteria for Olaparib and ATMi monotherapy.

**S5A:** Contains the number and frequency of patients with damaging BRCA1 or BRCA2 and ATM mutations in the indicated 30 PanCancer TCGA projects. To estimate the size of the benefiting patient population from this combination, we also involved cancer type prevalence data from the NCI- Surveillance, Epidemiology, and End Results Program by using the SEER explorer application. 109 out of 10141 (1.075%) patients met these conditions involved in 30 PanCancer TCGA studies. Our estimation indicates that more than 120,000 patients with various tumor types could benefit from Olaparib therapy combined with ATM inhibitors.

**S5B1:** Contains the number and frequency of patients bearing damaging mutations of the indicated genes in patients with the same tumor types, which we examined in *in silico* screens through cell lines. Mutational combination specific biomarkers were represented with low frequency across all indications that we investigated during *in silico* screens. Functional alteration of TP53BP1 were the most frequent in patients with cutaneous melanoma and colorectal cancers (6,3% and 3,7%, respectively), DDB1 functional alteration was also observed in melanoma and gastric cancer samples with ~2% frequency, while alteration of CUL4A were the most occurrent in melanoma patients. Mutational frequency distribution was determined by using the TCGA dataset. The extended mutational data was obtained from cBioPortal for each prioritized tumor types. The damaging status of the mutations was extracted from the PolyPhen feature. The final frequency table indicates the number of damaging mutations at combination biomarker and TCGA tumor type levels.

**S5B2:** Contains the number and frequency of patients, who express the given genes differently compared to the mean expression of total patients involved in TCGA studies. Like S5B1, we analyzed the data only those studies, which represents tumor types we modelled during *in silico* screenings by cell lines. Using gene expression z-score data from cBioPortal we examined the gene expression distributions of the potential expression biomarkers for each prioritized TCGA tumor types. For this, the *data_RNA_Seq_v2_mRNA_median_all_sample_Zscores* data was used which profiles the log-transformed mRNA expression z-scores compared to the expression distribution of all samples. In each tumor types we kept patients where the z-score absolute value was higher than 2 and then quantified these cases in each prioritized tumor types. We detected RAD54B/L overexpression significantly decreasing synergy. Interestingly, underexpression of these genes were the most prevalent with 51-54% frequency in prostate adenocarcinoma, kidney renal clear cell (KIRC) and bladder carcinomas (BLCA). The underexpression of ATRIP was the most prevalent in KIRC (54%) and squamous cell lung carcinomas (50,9%). KAT5 overexpression was observed in acute lymphoid leukemias (4%) and BLCA (3,9%).

Column legends can be found in the corresponding data files. Abbreviations for TCGA projects for better understanding can be found here: https://gdc.cancer.gov/resources-tcga-users/tcga-code-tables/tcga-study-abbreviations
